# An atlas of eukaryotic centromere architecture reveals recurrent evolutionary dynamics

**DOI:** 10.64898/2026.09.17.752431

**Authors:** Piotr Włodzimierz, Estela Perez-Roman, Jacob Gonzalez Isa, Michael Hong, Meng Zhang, Ludmila Oliveira, Katharine Jenike, Robin Burns, Pio Sierra, Matthias Heuberger, Chenxi Zhou, Sagarika Koppera, Mia Becker, Nicola Gorringe, Petr Novak, Yennifer Mata-Sucre, Giulio Formenti, Felipe Karam Teixeira, Ines Anna Drinnenberg, Ioannis Kontoyiannis, Kamil Jaron, Marcela Uilano-Silva, André Marques, Jiří Macas, Mark Blaxter, Richard Durbin, Tree of Life Consortium, Alexandros Bousios, Ian R. Henderson

**Affiliations:** Department of Plant Sciences, Downing Street, University of Cambridge, Cambridge, United Kingdom; Institute of Biochemistry and Biophysics, Polish Academy of Sciences, Warsaw, Poland; School of Life Sciences, University of Sussex, Brighton, United Kingdom; Department of Genetics, University of Cambridge, Cambridge, United Kingdom; Biology Centre, Czech Academy of Sciences, České Budějovice, Czech Republic; Department of Chromosome Biology, Max Planck Institute for Plant Breeding Research, Cologne, Germany; The Vertebrate Genome Laboratory, The Rockefeller University, New York, NY 10065, USA; Department of Physiology, Development and Neuroscience, University of Cambridge, Cambridge, United Kingdom; Institut Curie, PSL Research University, Sorbonne Université, CNRS, Paris, France; Statistical Laboratory, University of Cambridge, Cambridge, United Kingdom; Tree of Life, Wellcome Sanger Institute, Cambridge, United Kingdom; Faculty of Biosciences and Aquaculture, Nord University, Bodø, Norway

**Keywords:** Centromere, evolution, satellites, transposons, CENP-A, CENH3, holocentric

## Abstract

Centromeres evolved at the root of eukaryotes to segregate chromosomes during cell division^1–4^. Despite their ancient origin, centromeric DNA sequences evolve rapidly and adopt diverse architectures, including point centromeres, satellite arrays, transposon clusters, and holocentrics^2,5–12^. To analyse centromere evolution at a broad scale, we characterised architectures across 325 diverse Darwin Tree of Life genome assemblies^13^. Centromere architecture is evolutionarily labile, and similar configurations arise independently across divergent lineages. In plants and animals, we modelled centromere evolution as a recurrent cycle, in which satellite- and transposon-based architectures interconvert, with independent origins of holocentricity. We curated >23 million satellite repeats comprising 263 families from 165 species. Despite sequence divergence between satellite families, higher order repeats are prevalent, indicating constraint on repeat architecture rather than primary sequence. Satellite arrays are heavily invaded by diverse transposon families, consistent with convergent adaptation to the centromeric niche. In 89 species, transposons themselves constitute the primary centromere structure. We observed centrophilic transposons forming tandem arrays, suggesting mechanisms for satellite regeneration. Our sample includes five independent origins of holocentricity in plants and animals, which vary in association with periodic satellite arrays. We propose that genetic instability, centrophilic transposition, and transmission distortion promote recurrent centromere architectural interconversions during evolution.

## Introduction

Centromeres are essential chromosomal regions that load kinetochore complexes, enabling spindle microtubule attachment and accurate segregation during cell division^14,15^. Kinetochores and centromeres represent synapomorphies, likely arising early during eukaryogenesis ∼1.8-2.7 billion years ago, which are distinct from bacterial chromosome segregation systems, such as ParABS^1–3,16,17^. The deposition of nucleosomes containing the CENH3/CENP-A histone H3 variant typically determines where the kinetochore assembles^14,15,18–21^. The presence of a microtubule spindle, kinetochores, and CENH3/CENP-A are nearly universal across eukaryotes, suggesting these features were present in the Last Eukaryotic Common Ancestor (LECA)^3,17,22,23^; although derived exceptions exist including loss of CENH3 in holocentric insects and *Mucor* fungi^24,25^, and non-orthologous kinetochores in trypanosomes^26^.

Despite the ancient origin of centromeres and functional conservation, the underlying DNA sequences evolve rapidly and show high variability within and between species^1,2,5–11^. Centromere architectures range from compact point centromeres (∼120 base pairs in budding yeast)^11,27–29^, to megabase-scale regional centromeres built from satellite DNA arrays and transposon-rich sequences^7,11,12,30–33^, while other lineages adopt holocentric configurations with kinetochore assembly sites distributed along the chromosome length^24,34–37^. The evolutionary forces driving centromeric architectural transitions, and whether particular centromere types are constrained to specific lineages, remain unclear. Here, we explored the phylogenetic distribution of centromere architectures to test whether they follow predictable evolutionary dynamics.

### An atlas of centromeric architectures in 325 diverse eukaryotes

Recent advances in long-read sequencing technology and genome assembly algorithms enable repetitive centromere regions to be assembled at scale^7–9,12,13,31,35,36,38–42^. We used the Darwin Tree of Life (DToL) genome database to survey centromere architecture across 325 diverse eukaryotic species^13^ (**Supplementary Table 1**). This data source is uniquely suited for comparative centromere analysis due to its standardized assembly pipeline, extensive phylogenetic sampling, and chromosome-scale scaffolding with Hi-C data^13^. (**Supplementary Table 1**). The analysed genomes meet the Earth BioGenome Project 6.C.Q40 standard^38^, with megabase N50 contig continuity, chromosome-scale N50 scaffolding, and a less than 1:10,000 base error rate, making them suitable quality for mapping centromere architectures.

We designed our genome sampling to balance broad phylogenetic coverage with deeper representation in species-rich clades. Our sample focuses on Archaeplastida and Opisthokonta, which contain ∼90-95% of described eukaryotic species^43^. From the Archaeplastida, we selected 89 species; comprising 4 green algae, 11 non-vascular bryophytes, 31 monocotyledonous, and 43 dicotyledonous angiosperms. From the Opisthokonta, we selected 49 fungi, comprising 32 basidiomycetes, 15 ascomycetes, and 2 mucoromycetes; and from metazoa, we selected 184 genomes, including 30 chordates spanning fish, birds, non-avian reptiles, and mammals; as well as 126 arthropods that were predominantly insects across 16 orders, with an emphasis on Hymenoptera, Coleoptera, Lepidoptera, and Diptera, and 28 additional invertebrates representing 8 phyla. We included representatives from TSAR (2 alveolates) and Discoba (1 species), although several eukaryotic supergroups, for example Amoebozoa, Haptista, Cryptista, and Metamonada are unrepresented, reflecting a lack of chromosome-scale assemblies for these lineages in the DToL database. The atlas contains 12 groups of congeneric species, allowing local evolutionary comparisons in those cases. The 325 genomes span an ∼850-fold size range between 13.6 megabases (Mb) and 11.5 gigabases (Gb), across 2 to 67 chromosome pairs (**Supplementary Table 1**).

The rapid rate of centromere evolution means no single sequence can be used to identify these regions across eukaryotes^2,5–12^. However, satellites, defined as tandem repeats with monomers typically in the size range 100-200 base pairs (bp)^44^, and transposable elements, are established to be centromere-enriched, whereas protein-coding genes are rare^1,2,12,30–33^. Therefore, for each genome, we performed three complementary *ab initio* annotation approaches: gene prediction using Helixer, transposable element annotation using EDTA with lineage-specific RepBase libraries and TEsorter, and tandem repeat identification using TRASH^45–48^. We additionally quantified sequence entropy rate using the context-tree weighting algorithm^49,50^, which provides an annotation-free estimate of sequence predictability and repetition along chromosomes. We integrated these annotations into a Centromere Annotation Pipeline (CAP) that identifies candidate centromeres by searching for regions of (i) low protein-coding gene density, (ii) satellite repeat and/or transposon enrichment, and (iii) low sequence entropy rates (**Extended Data Fig. 1**).

As satellite array architecture is common in eukaryote centromeres^1,2,7,30,31,39,40,51,52^, CAP first scores tandem repeats. Satellite families are scored more highly if they form, (i) longer, contiguous arrays, (ii) are shared on multiple chromosomes, (iii) show high sequence homogenization consistent with concerted evolution^51,53^, and are located in (iv) gene-poor, and (v) transposon-enriched regions. If multiple high-scoring satellite families are present, the genome is classified as ditypic (two families), or polytypic (three or more families). If satellite repeats are not identified, CAP searches for regions of transposon-enrichment that are shared between chromosomes, as candidate transposon-based architectures. For the 41 species from known holocentric taxa, we used CAP to search for periodically distributed short satellite arrays spread along the chromosomes, which are characteristic of holocentric plant species^35,36,40^, and *Meloidogyne* nematodes^54^. In fungal genomes, centromeric and pericentromeric inter-chromosome Hi-C contacts can provide orthogonal evidence for centromere locations^55,56^, which we used together with CAP.

To experimentally validate centromere predictions, we performed fluorescent *in situ* hybridization (FISH) with candidate satellite probes and immunostaining for the conserved kinetochore protein KNL1^57^ in four plants; *Chamaenerion angustifolium* (Myrtales), *Mercurialis annua* (Malpighiales), *Linaria vulgaris*, and *Ballota nigra* (Lamiales). We used KNL1 as a kinetochore marker, due to antibody cross-reactivity across diverse plant species^57^. In each species, we confirmed monocentric KNL1 localization, and that the CAP-predicted satellites localized to chromosome centromere constrictions (**Extended Data Fig. 2a-2l**). In *M. annua*, the chromosomes contain megabase tandem repeat arrays built from distinct 60 and 99 bp-long satellites (**Extended Data Fig. 2d**). The 99 bp satellites were assigned a higher score due to their greater homogeneity; and FISH confirmed that they, rather than the 60 bp family, localized to centromere constrictions (**Extended Data Fig. 2f**).

To validate transposon-dominated centromere architecture, we used FISH to detect candidate Athila retrotransposons identified by CAP in the plant *Geum urbanum* (Rosales), along with KNL1 immunostaining. We confirmed monocentric localization of KNL1, and Athila FISH staining in centromeric constrictions (**Extended Data Fig. 2m-2o**). We cytogenetically confirmed the centromeric location of CAP-predicted Gypsy-CRM and Gypsy-Tekay retrotransposons in *Ailanthus altissima* (Sapindales) and *Solanum dulcamara* (Solanales), respectively (**Extended Data Fig. 2p-2u**). Beyond monocentric architectures, we validated predictions in the holocentric species *Luzula sylvatica* (Poales), where multiple kinetochore and CENH3 loading sites occur on regularly spaced arrays of 125 and 174 bp satellite repeat arrays^40^ (**Extended Data Fig. 2v-2x**), which were correctly identified by CAP.

Finally, we validated CAP against 8 plant, fungal, and metazoan genomes where centromeric chromatin has been previously mapped via ChIP-seq of CENH3/CENP-A^12,31,33,39,58–60^ (**Extended Data Fig. 3**). CAP identified satellite repeats as candidate centromeres in human, mouse, rice, maize, and pea, as well as candidate transposon centromeres in wheat and *Cryptococcus*, which are known to be CENP-A/CENH3-occupied^12,31,33,39,58–61^ (**Extended Data Fig. 3**). In the stick insect *Timema*, CAP classified a cryptic centromere state, consistent with the CENP-A occupied regions lacking shared repeats^61^ (**Extended Data Fig. 3**). Therefore, our workflow reliably identifies known monocentric satellite and transposon centromere architectures, as well as periodic satellite arrays associated with holocentric species. In species where the centromeres are not consistently associated with DNA repeats^27–29,34,62,63^, CAP outputs an unknown (cryptic) state.

### A recurrent model for centromere evolution across eukaryotes

Using CAP, we assigned 288 (89%) genomes to a centromere architectural class. 147 (45%) species were classified as satellite-type (which we term α-state), 89 (27%) as transposon-type (which we term Ω-state), and 11 (3%) showed mixed α/Ω states on different chromosomes (**Fig. 1a** and **Supplementary Table 1**). 41 (13%) species were from known holocentric clades, 19 (6%) of which had periodic satellite repeats. In 6 (2%) species, Hi-C interchromosomal contacts identified centromere regions, but these did not show an associated repeat sequence enrichment. In 31 (10%) species, CAP detected neither centromeric satellites, nor transposons, and these species do not belong to known holocentric taxa. These architecturally cryptic genomes represent a biologically significant class and may contain previously unrecognised repeatless holocentric species^24,34,37^, or non-repetitive centromeres, analogous to those in budding yeast and horses^27–29,62^.

**Figure 1.**
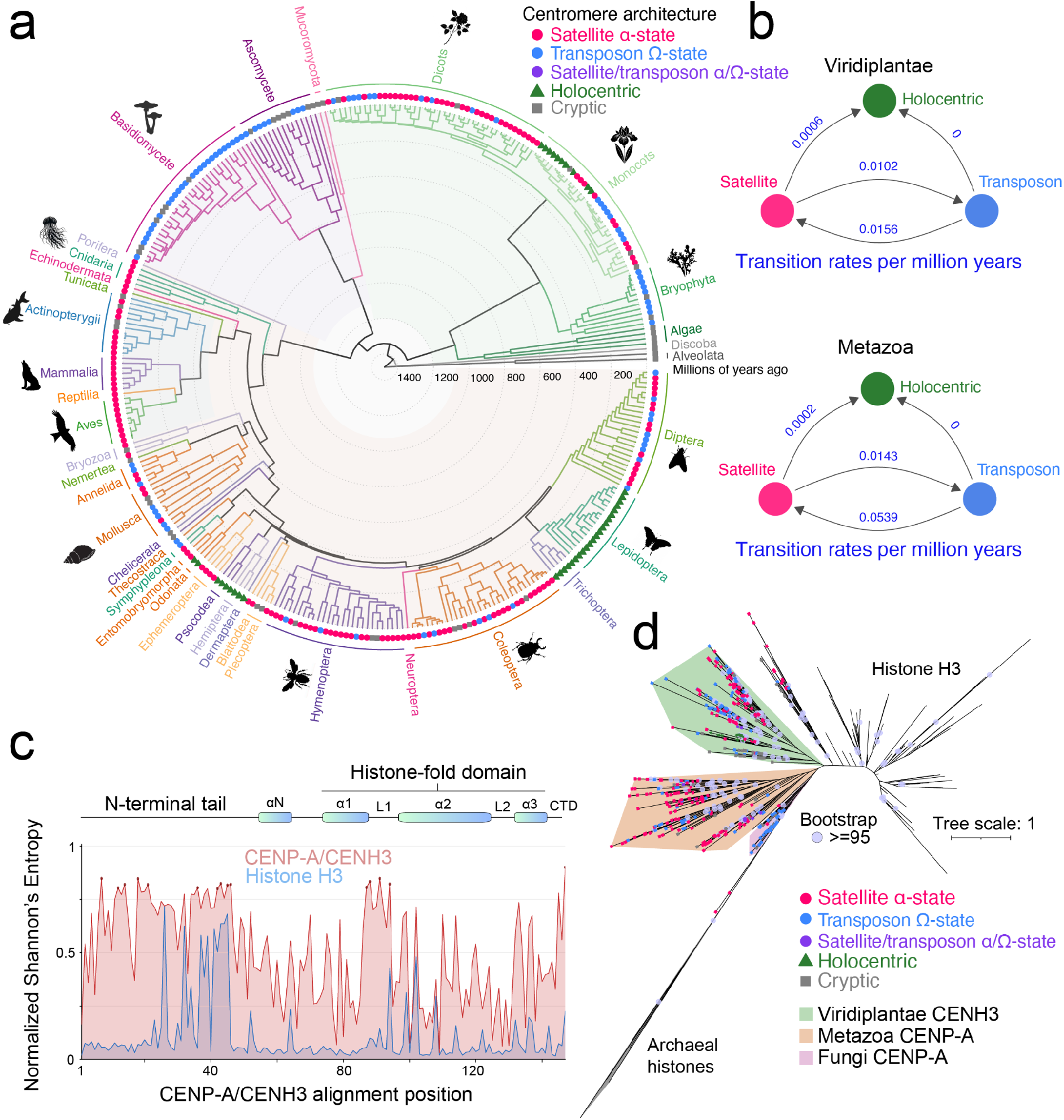
An atlas of eukaryotic centromere architecture reveals a recurrent model of evolution. **a.** Phylogenetic species tree generated using a BUSCO gene multiple sequence alignment using FastSpeciesTree^64^, with branch lengths in millions of years (Mya) from TimeTree^67^ fossil calibration. CAP-derived centromere architecture classifications are indicated on branch tips as α-state (Satellite, pink), Ω-state (Transposon, blue), mixed α-satellite/Ω-transposon (purple), Holocentric (green), or Cryptic (grey). **b.** Flow diagrams of centromere architecture transitions within Viridiplantae (upper) and Metazoa (lower) based on ancestral state reconstruction using the ARD_irrevH model in phytools^68^. Centromere architecture transition rates per million years from a time-calibrated chronogram are labelled in blue. **c.** Positional Shannon’s entropy of CENP-A/CENH3 (red) and H3-like (blue) sequences across a trimmed alignment computed from tips of the gene tree. Entropy is normalised to values between 0 and 1. Positions where the within-group gap frequency exceeds 85% were masked. Shaded dots mark the 10% highest-entropy CENP-A positions. Coloured bars at the top of the plot indicate predicted histone α-helical regions of CENP-A mapped from an AlphaFold2 structural model (UniProt Q8RVQ9) using STRIDE^78^, and projected onto the trimmed alignment coordinates. **d.** Phylogenetic tree of CENP-A/CENH3, canonical histone H3, and archaeal histone sequences. Tree nodes with bootstrap support >=95% are indicated by grey dots. Within the CENP-A/CENH3 clade, the centromere architectural state of the originating genome is indicated on the tree tips, as shown in a. Green, orange, and purple shading indicate clades of CENH3/CENP-A sequences from Viridiplantae, Metazoa, and Fungi, respectively.

To build a phylogenetic species tree, we used FastSpeciesTree^64^, which identifies conserved BUSCO genes directly from input proteomes via DIAMOND, reconstructs pseudo-alignments from the high-scoring segment pairs, and concatenates these into a supermatrix for tree inference. A maximum-likelihood species tree was then inferred with IQ-TREE using ModelFinder^65^. The tree was dated using chronos^66^, and 62 time-calibration points from TimeTree^67^ (**Fig. 1a** and **Extended Data Fig. 4**). When CAP centromere architectures were displayed on the species tree, we observed phylogenetic dispersion of states, where closely related species frequently differed, and distantly related lineages converged on similar architectures (**Fig. 1a**). To model centromere evolution, we applied ancestral state reconstruction with phytools^68^, using the whole tree, or sub-trees from two major clades; Metazoa and Viridiplantae (**Fig. 1b** and **Extended Data Fig. 5**). Comparisons favoured a model in which satellite and transposon architectures transition under all-rates-different (ARD) constraints, and holocentrics are an evolutionary sink (**Fig. 1b** and **Extended Data Fig. 5**). We observed clades enriched for specific centromere architectures, including satellite α-states in vertebrates (25/29), dicots (31/43), and coleoptera (21/30); and transposon cluster Ω-states in bryophytes (9/11), fungi (34/49), and diptera (10/24). We note that cryptic centromere states were present in all protists (3/3) and algae (4/4) sampled. This supports a recurrent model for centromere architecture evolution within the Viridiplantae and Metazoa, together with clade-specific biases that indicate varying evolutionary equilibria in different lineages.

To investigate how protein-level evolution relates to the architectural cycles observed in centromeric DNA, we examined variation in histone CENH3/CENP-A and compared to canonical histone H3. We retrieved 422 CENH3/CENP-A orthologs from the 325 genomes, along with 897 canonical histone H3-like proteins, as well as histone-related proteins from archaeal Asgard, Euryarchaeota, and DPANN lineages as an outgroup, and constructed a phylogenetic tree (**Fig. 1d** and **Extended Data Fig. 6a**). We observed longer terminal branch lengths within the CENH3/CENP-A clade relative to histone H3 (Wilcoxon test *P*<0.001) (**Fig. 1d** and **Extended Data Fig. 6b**), which is consistent with accelerated evolution^18,69^. Positional Shannon’s entropy analysis of CENH3/CENP-A alignments revealed that sequence variability was non-uniformly distributed across the protein. The core histone fold domain (HFD) α-helices and loop 2 show relatively low entropy (**Fig. 1c** and **Extended Data Fig. 6a**), likely reflecting structural constraints relating to nucleosome function. In contrast, the CENH3/CENP-A N-terminal tail, the first loop of the HFD, and the C-terminal region show elevated entropy across all taxonomic groups (**Fig. 1c** and **Extended Data Fig. 6a**). This is consistent with adaptive evolution in CENH3/CENP-A regions known to be involved in kinetochore interactions and centromere targeting^70–74^. Mapping centromere architecture onto the histone phylogeny revealed that satellite, transposon, and holocentric types were broadly distributed across the CENH3/CENP-A clade (**Fig. 1d**). To test for residues that are diagnostic of centromere architecture class, we used GroupSim^75^, which identifies positions conserved within groups of interest, but divergent between them. As a positive control, we compared CENP-A/CENH3 and H3 groups, which recovered known sequence differences^76,77^ (**Extended Data Fig. 6c**). When comparing α-state and Ω-state centromere architecture classes, no group-specifying CENH3/CENP-A positions were identified (**Extended Data Fig. 6d**). Together, this indicates that although CENH3/CENP-A evolves more rapidly than canonical histone H3, its sequence divergence is uncoupled from underlying centromere DNA architectural states at this phylogenetic scale.

### Satellite repeat diversity and recurrent higher order structures

At one pole of the inferred centromere evolutionary cycle, architectures are dominated by satellite repeats (α-state) (**Fig. 1b**). Across species in which we identified satellite-type centromere architectures, in addition to the holocentric species where we found periodic satellite arrays, we curated a total of 23,577,177 repeats, representing 263 distinct families (**Fig. 2a** and **Supplementary Table 2**). Of these genomes, 118 were monotypic, 27 were ditypic, and 20 were polytypic, with respect to the number of satellite families CAP identified as centromere candidates (**Supplementary Table 2**). Each satellite family was named using the species Tree of Life Identifier (ToLID) followed by the mean monomer length. For instance, the ∼165 bp satellite family in the beetle *Ocypus olens* is designated icOcyOlen.165.

**Figure 2.**
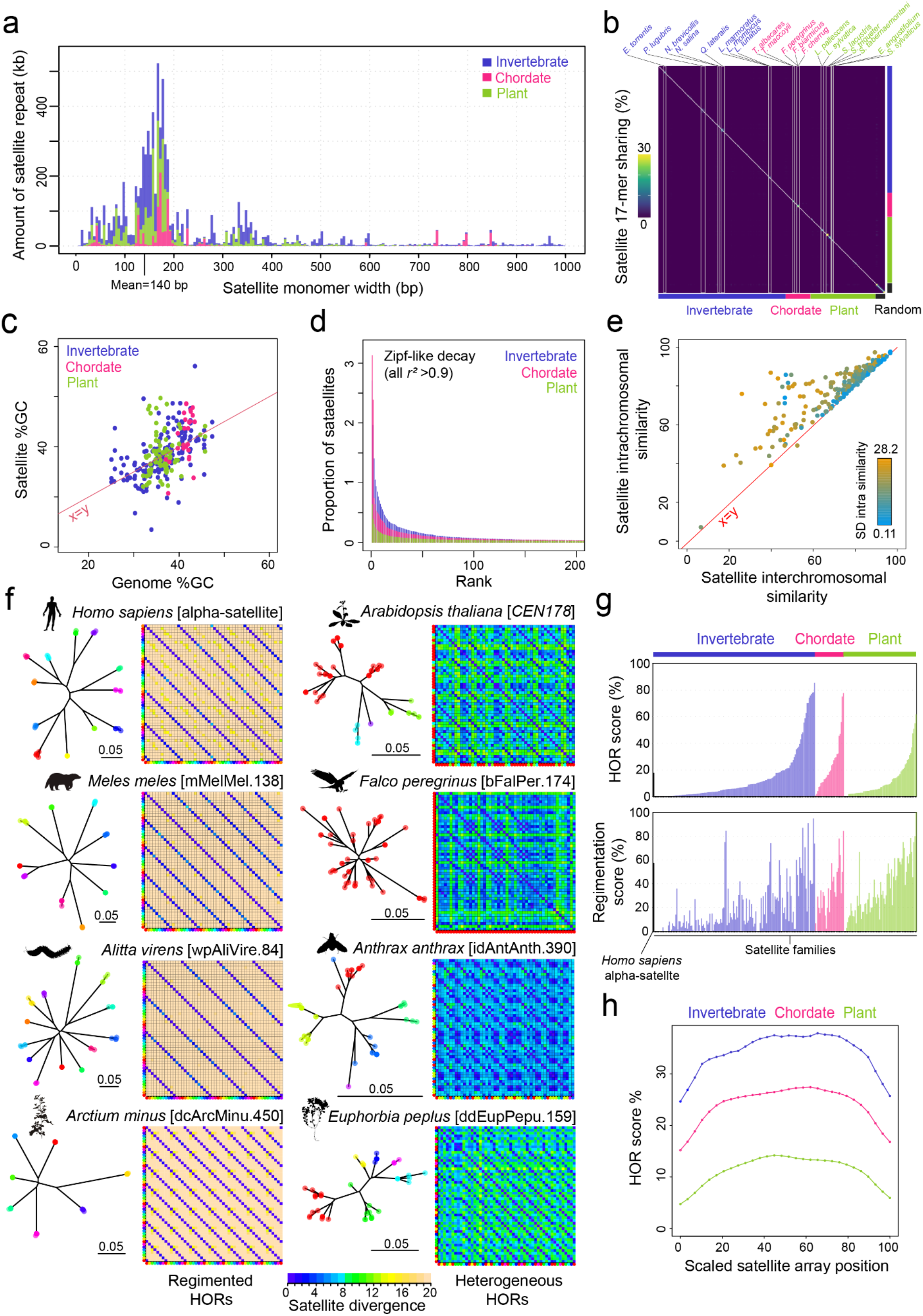
Satellite diversity and recurrent higher order repeats. **a.** Histogram of the amount of satellite sequence (kb) at varying monomer widths (base pairs), coloured according to plant (green), chordate (pink), or invertebrate (blue) taxa. Up to 50 kb of repeats were randomly selected per family. The mean satellite length of 140 bp is labelled. **b.** Heat map of 17-mer Jaccard sharing between satellite families from plants, chordates, and invertebrates, and random sequences with varying GC content, with samples ordered phylogenetically. White boxes highlight satellites within the same genome, or in closely related species, that show sequence similarity above background levels. **c.** Scatterplot of %GC for satellite repeat families, compared to %GC of the cognate genome, coloured by plants (green), chordates (pink), or invertebrates (blue). The red diagonal line indicates a linear x=y relationship. **d.** For each satellite family, copies were ranked by abundance and their proportional frequency plotted, separately for plants, chordates and invertebrates. When transformed in a log-log analysis these data conform to a Zipf’s type decay (all *r*^2^>0.9). **e.** Scatterplot for each satellite repeat family showing mean inter-chromosomal similarity compared to mean intra-chromosomal similarity. The red diagonal line indicates the linear x=y relationship. Points are coloured by the standard deviation in satellite intrachromosomal similarity values, according to the inset scale. **f.** For each species, a sample of 50 contiguous satellite repeats were analysed and their pairwise divergence plotted in a 2D matrix. A phylogenetic tree of the same repeats is shown to the left, where clade tips are coloured, and the same colours mark the same repeats displayed on the x and y axis of the divergence matrix shown to the right. The examples on the left show regimented HORs like humans, whereas those on the right show heterogeneous HORs like Arabidopsis. **g.** Barplots quantifying higher order repeats (Upper, %) and HOR regimentation (Lower, %) in invertebrate, chordate, and plant satellite arrays. On the left of the plot, we include analysis of regimented alpha-satellite arrays in *Homo sapiens* genome assembly CHM13^42^. HOR score quantifies repeat participation in higher order structures, and the regimentation score quantifies the frequency of periodic HORs. **h.** For satellite arrays from invertebrates, chordates, and plants, we quantified HOR score (%) with respect to the scaled length of the arrays. To analyse HOR scores along the scaled length of centromeres, each array was divided into 30 length-scaled windows. Within each window, the summed HOR score was divided by the number of repeats in that window.

The satellites were enriched in the 100-200 bp range (128/263 families; overall mean=140 bp), consistent with each repeat wrapping a single nucleosome^44,51,79^ (**Fig. 2a** and **Supplementary Table 2**). However, substantial sub-nucleosomal (<100 bp) satellites were identified (32 families), alongside supra-nucleosomal repeats in the 201 bp to 5 kb range (103 families). Given the diversity in monomer lengths, we asked whether satellite families shared sequence identity. We converted satellites from each family into k-mers of length 17 and calculated pairwise Jaccard similarity indices, benchmarking against randomly generated sequences (**Fig. 2b**). Overall satellite similarity was low (average sharing of 17-mers=0.03%), although higher levels were observed between repeats of congeneric species. For example, three *Falco* genomes contain related ∼174 bp satellites with 15-26% shared 17-mers (**Fig. 2b**). A subset of ditypic and polytypic families found within the same genome shared 2-11% of 17-mers, suggesting shared ancestry (**Fig. 2b**). Satellite families retained similarity (>80% identity) up to ∼26 million years (My) of divergence, with a mean similarity decay half-life of ∼12 My (**Extended Data Fig. 7a**). This confirms that satellite sequence turnover is rapid, with homology erased over relatively short evolutionary timescales^44,51,79^. The absence of satellite sequence conservation extends to base composition. Although human satellite and yeast centromeres are AT-rich relative to their genome^28,31^, this pattern did not generalise. Across the satellite families, 49% (128 of 263) were less AT-rich than their genome average, while 51% (135 of 263) were more AT-rich (**Fig. 2c**). The absence of conservation and compositional bias shows that primary sequence is not a constrained feature of centromere satellites at this scale.

As satellite sequences were not conserved across families, we explored whether commonalities exist at the level of repeat organisation. We analysed satellite repeat frequency distribution within each family, which strongly conform to Zipf’s law (frequency∝1/rank; *r²*>0.9 log-log regression, across all families) (**Fig. 2d**). This is consistent with copy number-dependent amplification pathways, such as unequal homologous recombination, in which variants amplify at rates proportional to their abundance^44,51,80^. Within-chromosome satellite similarity was nearly always greater than between-chromosome similarity (**Fig. 2e**, points above the *x*=*y* diagonal), consistent with intra-chromosomal recombination operating more frequently than inter-chromosomal exchange. The standard deviation of intra-chromosomal satellite similarity also increases with the difference between intra- and inter-chromosomal similarity (Spearman’s *ρ*=0.70 *P*<0.001) (**Fig. 2e**, colour scale). This supports that satellite families tend to follow independent evolutionary trajectories on each chromosome over time, consistent with recombination driving accumulation of chromosome-specific variation.

Satellite higher order repeats (HORs) reflect ongoing homogenization of array sequence, via homologous recombination^44,51,79^. For example, human centromeres show regimented HOR structures with periodicities ranging between 2 to 46 alpha-satellite monomers, over kilobase to megabase scales^31,46,51,81–83^. In contrast, although the plant *Arabidopsis thaliana* contains abundant centromeric satellite HORs, they show heterogeneous spacing and lack the long-range periodicity observed in humans^8,30,46^ (**Fig. 2f**). To investigate the extent of regimented versus heterogeneous HORs across the DToL satellites, we used TRASH to perform all pairwise comparisons of satellite monomers within each centromeric array^46^, identifying paired blocks of contiguous monomers with similarity above a defined threshold (**Fig. 2f** and **Extended Data Fig. 8**). Higher order satellite repeat structure was prevalent, and we observed plant and animal genomes that contained regimented and heterogeneous HORs, although satellite families varied in the proportion of each type of organisation (**Fig. 2f-2g** and **Extended Data Fig. 8 and 9**). We quantified the spatial distribution of HOR blocks within the arrays and observed that frequency was highest in the array centres (**Fig. 2h**). This pattern is consistent with internal satellite repeats having more opportunities for recombination than those located near the array edges^7,51,81^.

While most satellite-state genomes were classified as monotypic (n=118), 27 were ditypic, and 20 were polytypic (**Supplementary Table 2**). These genomes varied in the extent to which satellite families shared chromosomes. For example, ditypic satellites can be intermixed within the same centromere region (e.g. icScaQuad.92/icScaQuad.109 and idCluTigr.118/idCluTigr.193), or families can be segregated on alternate chromosomes (e.g. dcPolTetr.136/dcPolTetr.449) (**Extended Data Fig. 7b**). In *Abia candens* (Hymenoptera), each acrocentric chromosome possesses a tiered arrangement of a sub-telomeric iyAbiCand.146 array juxtaposed with an internal iyAbiCand.143 array (**Extended Data Fig. 7b**). The ditypic and polytypic genomes may reveal snapshots of potential competitive or co-operative dynamics between evolving satellite families. Other species show clustering of non-contiguous satellite arrays, such as in *Ricordia florida* (Cnidaria) and *Misopates orontium* (Angiosperm) (**Extended Data Fig. 7c**), which are reminiscent of metapolycentric genomes such as pea where an extended primary constriction contains multiple centromeric satellite arrays^39,84^ (**Extended Data Fig. 3g**).

### Convergent centrophilic transposon adaptation across eukaryotes

Within the inferred centromere evolutionary cycle, α-state satellite repeats become progressively invaded by centrophilic transposons, ultimately entering the Ω-transposon state (**Fig. 1b**). To investigate the frequency and diversity of centrophilic transposon invasions, we extracted non-satellite ‘interruptions’ (<100 kb) from the satellite centromeric arrays and quantified their transposon content (**Fig. 3a**). On average, the interruptions comprise 17% of centromere array space, of which 70% was annotated as transposon (**Fig. 3a**). We observed varying degrees of transposon invasion, where the plants had the highest array transposon content (18%), followed by invertebrates (10%), and chordates (7%) (**Fig. 3a**). At one extreme, species such as *Quercus robur* (Fagales) and *Taurulus bubalis* (Perciformes) contain near-pristine satellite arrays with minimal transposon content (0.34% and 0.04%, respectively), while at the other, *Coccinella septempunctata* (Coleoptera) and *Nomada fabriciana* (Hymenoptera) harbour satellite arrays that are heavily fragmented by transposons (25% and 18% transposons, respectively) (**Fig. 3a-3b**). We propose this reflects satellite arrays relatively close to the α-state, versus those relatively close to the Ω-state, respectively.

**Figure 3.**
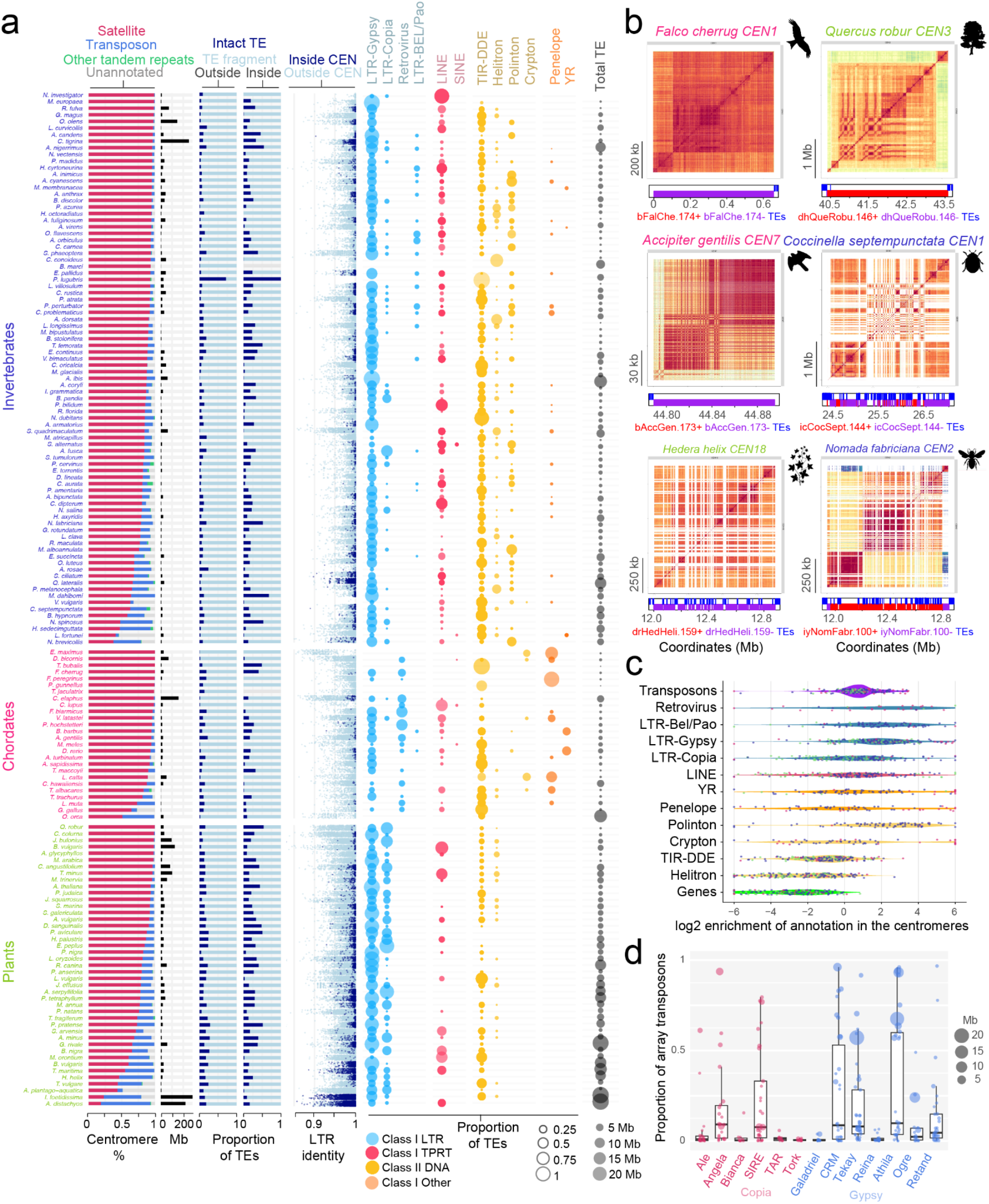
Convergent centrophilic adaptation of diverse transposon lineages across eukaryotes. **a.** For genomes classified with α-state satellite architecture, we defined centromeric arrays and their non-satellite interruptions and quantified (i) the proportion of array sequence that overlapped satellites (pink), transposons (blue), other tandem repeats (green), or unannotated (grey) sequence, (ii) the total centromere array size in megabases (Mb), (iii) the proportion of transposon annotation that was ‘intact’ (dark blue) versus ‘fragmented’ (light blue) inside versus outside the centromere satellite arrays, (iv) for intact LTR elements inside (dark blue) and outside (light blue) the satellite arrays, we plotted the paired LTR identity, and, (v) coloured bubbles represent different classes of transposable element proportional to their abundance within the centromeric arrays, (vi) total transposon space within the arrays, capped at 20 Mb. **b.** ModDotPlot^88^ sequence identity heat maps are shown from representative satellite centromeres, where red shading indicates the highest levels of sequence identity. Beneath each heatmap, satellite forward (red) and reverse (purple) strand, and transposon (blue) annotation are indicated by coloured bars. To the left of each heatmap is a scale bar showing physical distance in kb or Mb. The plots are arranged by the lowest degree of transposon invasion (top), to the highest (bottom). **c.** Log_2_ enrichment of the listed sequence annotations found in the non-satellite interruptions of arrays, compared to the rest of the genome. Each data point represents a species with violin plots behind showing the data distribution. Points are coloured according to origin in a plant, chordate, or invertebrate genome. **d.** Plots showing the relative proportion and amount (Mb indicated by circle size, key to right) of LTR retrotransposon lineages found within centromere satellite arrays of α-state plant species (each data point is a species).

Class I LTR-retrotransposons showed strongest enrichment in the satellite arrays across taxa (11% of centromeric sequence across all species) (**Fig. 3a** and **3c**). Within the LTR order, Gypsy is the most enriched superfamily, followed by Copia; with retroviruses frequent in the chordates, and Bel-Pao in the Metazoa (**Fig. 3a** and **3c**). LINE non-LTR retrotransposons are found in centromere arrays across taxa (1.3% of centromeric sequence) (**Fig. 3a**). Although retrotransposons overall were dominant in the satellites, we observed substantial Class II DNA transposons (2%), including DDE-integrase class terminal inverted repeat transposons (TIRs), and rolling-circle Helitrons (**Fig. 3a**). Less frequent, but notable, were tyrosine recombinase and Penelope transposons in chordates (**Fig. 3a**). In plants, lineage-resolved classification of LTR retrotransposons^85^ allowed us to observe multiple satellite colonizations by diverse lineages; most frequently by Gypsy CRM and Tekay chromoviruses, non-chromovirus Athila, and Angela and SIRE Copia lineages (**Fig. 3d** and **Extended Data Fig. 10**). Within Athila alone, phylogenetic analysis supports independent centromere colonizations in at least 13 angiosperms (**Extended Data Fig. 11**). As the centrophilic elements identified span the known mechanistic diversity of eukaryotic transposons, including DDE-integrase (LTR and TIR), target-primed reverse transcription (LINEs), tyrosine-recombinases, HUH-endonuclease rolling-circle replication (helitrons), and GIY-YIG-endonuclease-primed reverse transcription (Penelope), which diverged over 1-2 billion years ago^86,87^, this supports convergent adaptation of diverse transposon lineages to the satellite centromeric niche, across eukaryotes.

In satellite arrays that contained LTR retrotransposons, we quantified the proportion of structurally intact elements, defined as those with paired LTRs and internal coding sequences, versus fragmented copies, and their relative abundance inside versus outside of the satellite arrays (**Fig. 3a**). LTR elements located within arrays were significantly more structurally intact than those found outside centromeric regions (Welch Two Sample t-test, *P*<9.85×10^-7^) (**Fig. 3a**), consistent with recent insertion. To estimate transposable element age, we quantified sequence identity between the 5′ and 3′ long terminal repeats of intact LTR transposons, which are identical at the time of insertion, but diverge over time through mutation. Across all genomes, elements embedded in the satellite arrays had significantly higher LTR identity compared to those outside (Welch two sample t-test, *P*<1.55×10^-12^), indicating they are on average younger (**Fig. 3a**). These results are consistent with recent integration of centrophilic retrotransposons into satellite arrays, and rapid turnover across eukaryotes.

### Transposon-dominated centromeres in plants, animals, and fungi

In the Ω-state, we propose that a centrophilic transposon simultaneously targets centromeric chromatin during integration, as well as being the site of CENP-A/CENH3 enrichment itself; creating a transposon cluster on each chromosome. This predicts that Ω-state centromeres should consist of young, nested insertions of the centrophilic transposon family. An archetypal Ω-state is found in *Geum urbanum* (Rosales), where centromeres are composed almost entirely of Gypsy-Athila LTR retrotransposons (87.5% of ∼18 Mb centromere sequence) (**Fig. 4a-4i**). In contrast, the sister species *Geum rivale*, which diverged ∼2 million years ago^89^, shows an α-state satellite architecture at syntenic locations, with a dominant drGeuRiva.159 satellite repeat (65.7% of ∼57 Mb centromere sequence) (**Fig. 4b-4i** and **Extended Data Fig. 12**). *G. urbanum* centromeres contain a smaller, diverged population of drGeuRiva.159 copies compared to *G. rivale* (3,187 vs 236,781), while *G. rivale* maintains similar centromeric Athila content to *G. urbanum* (19.6 Mb vs 15.7 Mb, respectively) (**Fig. 4c-4g**). This is consistent with a transition between α-state and Ω-state centromere architecture within *Geum* that involved satellite expansion or contraction since speciation. In contrast, the other congeneric groups sampled in the atlas showed stable within-genus centromere architecture classifications (**Supplementary Table 1**).

**Figure 4.**
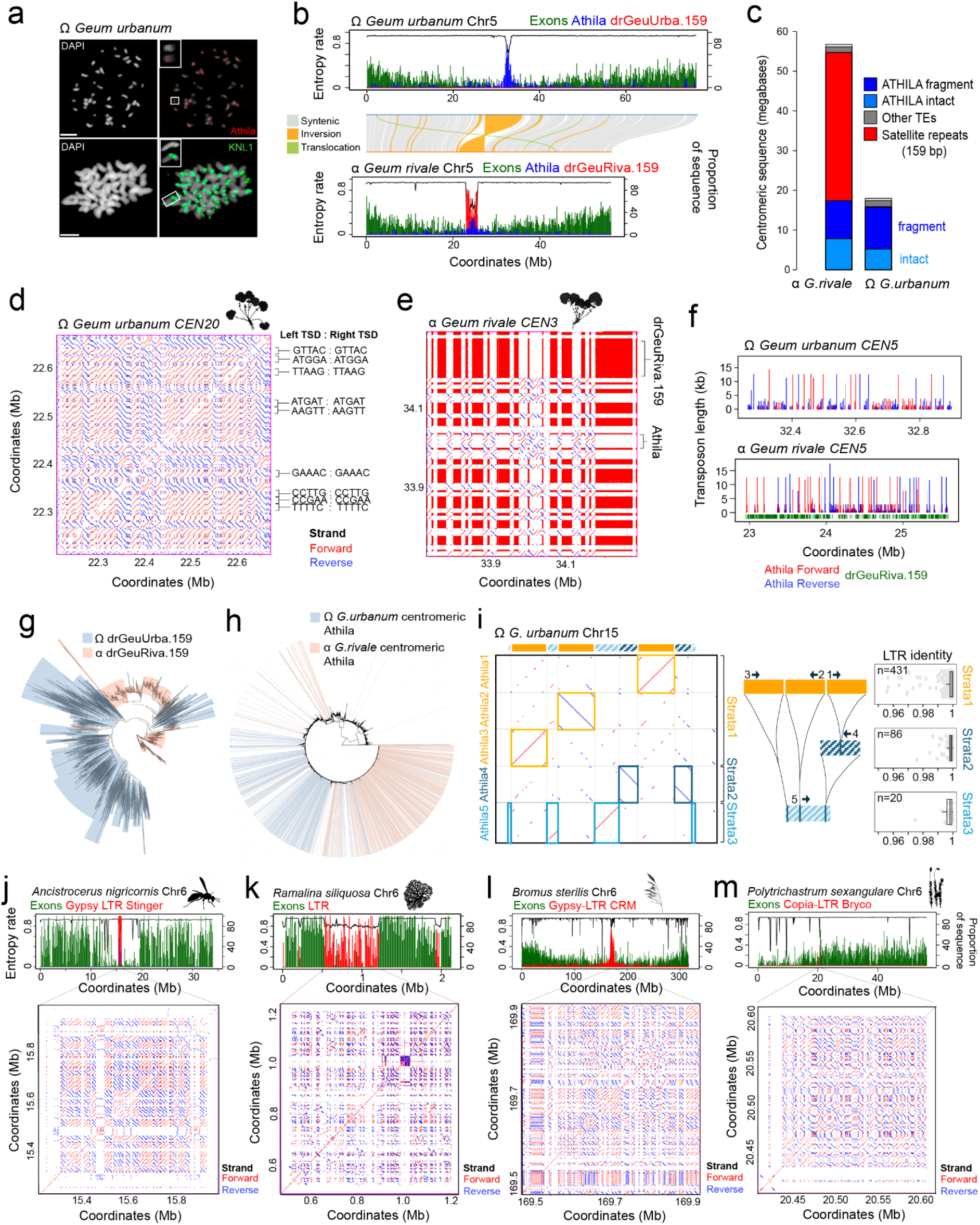
Transposon-dominated centromeres across eukaryotes. **a.** Cytogenetic analysis of *Geum urbanum*, with DNA stained by DAPI, immunostaining for KNL1 (green), and FISH staining for Athila centromeric transposons (red). Scale bars=5 µM. Reproduced in Extended Data Fig. 2. **b.** Plots of *G. urbanum* and *G. rivale* chromosome five, showing the proportion of sequence corresponding to gene exons (green), Athila retrotransposons (blue), and 159 base pair satellite repeats (red), and sequence entropy rate (black). The chromosomes are connected showing regions of synteny (grey), inversions (orange) and translocations (green) from SyRI^90^. **c.** Barplot showing *G. urbanum* and *G. rivale* centromere % composition by intact and fragmented Athila elements (blue), other transposons (dark grey), and 159 bp satellite repeats (red). **d.** A sequence identity dotplot within *G. urbanum CEN20*, with similarity on forward (red) and reverse (blue) strands displayed. For intact Athila elements within this region, their target site duplications (TSDs) are labelled and listed alongside. **e.** As for d, but showing a comparable region of *G. rivale CEN3*. Labels to the right indicate representative regions of Athila transposon and drGeuRiva.159 satellite sequences. **f.** Plots of *G. urbanum* and *G. rivale CEN5* showing the length of Athila annotations on forward (red) and reverse (blue) strands. In the *G. rivale* plot the position of drGeuRiva.159 repeats is also indicated along the x-axis (green). **g.** Phylogenetic tree of drGeuUrba.159 (blue) and drGeuRiva.159 (red) satellite repeats. **h.** Phylogenetic tree of intact Athila copies annotated genome-wide from *Geum urbanum* and *Geum rivale*. Shading indicates copies present in the centromeres. **i.** To the left, a dotplot of *G. urbanum* chromosome 15 along the x axis, compared against the contained intact Athila ordered by nesting strata; with forward and reverse strand similarity shown in red and blue, respectively. At the top, Athila elements are annotated in colours that match the nesting diagram shown to the right, with strata 1, 2, and 3 elements indicated, in addition to their nested integration positions (vertical lines). To the right are LTR identity barplots for Athila found genome-wide in the three nesting strata. **j.** Gene exons (green) and Gypsy-LTR (red) density along *Ancistrocerus nigricornis* chromosome 6, with a sequence identity dotplot beneath the centromere region, as in d. **k.** As for j, but showing chromosome 6 of the fungi *Ramalina siliquosa*. **l.** As for j, but showing chromosome 6 of the monocot *Bromus sterilis* and a LTR Gypsy-CRM centromere cluster. **m.** As for j, but showing chromosome 6 of the moss *Polytrichastrum sexangulare* and a LTR Copia-Bryco centromere cluster.

We identified (29.1% of total centromere sequence) and 758 (13.9%) intact centromeric Athila in *G. urbanum* and *G. rivale* respectively that form a single phylogenetic group (**Fig. 4h** and **Extended Data Fig. 12d-12e**), most of which are ∼12.5 kb in size and have high 5′-3′ LTR identity (133 and 276 have identical LTRs), indicating recent integration (**Fig. 4c** and **4f**). To assess whether *G. urbanum* centromeric Athila are nested, we computationally removed intact elements, patched their flanking sequences, and tested whether this reconstructed older strata of intact Athila (**Fig. 4i**). This approach demonstrated an extensive intact-within-intact nesting pattern, reconstituting composite elements in up to three strata, in total recovering 106 intact elements (**Fig. 4i**). The Athila in all strata were young, with median LTR identities >99.8%, consistent with recent transposition (**Fig. 4i**). Athila target site duplications (TSDs), which serve as fingerprints of independent integrations, were not widely shared within *G. urbanum* or *G. rivale* (e.g. **Fig. 4d**). The most frequent TSD (ATCAT) in *G. urbanum*, appeared only seven times, and never in tandem, supporting independent Athila insertions. The abundance of young, intact, and nested Athila elements, suggests that the *G. urbanum* centromeres are primarily shaped by ongoing centrophilic retrotransposition, which typifies the Ω-state, rather than post-integration amplification by tandem duplication.

We sought to identify Ω-state centromere architectures in other animal, plant, and fungal genomes that lacked satellite arrays. In the wasp *Ancistrocerus nigricornis*, each chromosome shows a central ∼200-600 kb gene-depleted region that contain clusters of Gypsy-LTR transposons, with approximately equal copies on either strand, and which show elevated inter-chromosomal Hi-C contacts (**Fig. 4j** and **Extended Data Fig. 13**). These elements are errantivirus endogenous retroviruses that we name Stinger (**Fig. 4j** and **Extended Data Fig. 13**). In the ascomycete Lichen-symbiote fungus *Ramalina siliquosa*, the chromosomes contain central 200-500 kb regions with depleted genes that are enriched for LTR transposons (**Fig. 4k**). The monocot *Bromus sterilis* contains 1-2 Mb deeply nested clusters of LTR-Gypsy CRM elements (**Fig. 4l**), whereas the moss *Polytrichastrum sexangulare* shows centromeric clusters of Bryco LTR-Copia elements (**Fig. 4m**). To compare the nesting depth of these arrays, we iteratively excised intact elements and re-annotated each layer, until no further elements were found. The *B. sterilis* centromeres had the deepest nesting layers (mean±1SD, 2.86±0.69), followed by *G. urbanum* (2.00±0.95), *G. rivale* (1.68±0.89), *P. sexangulare* (0.69±0.75), *R. siliquosa* (0.22±0.55) and *A. nigricornis* (0.17±0.41). Together, these genomes provide examples of transposon-based centromere architectures arising in diverged species from distinct transposon lineages.

### Formation of tandem repeats by centrophilic transposons

In the inferred centromeric evolutionary cycle, Ω-states can emerge from α-states through centrophilic transposon invasion (**Fig. 1b**). We next considered how transposon cluster Ω-states may return to satellite-dominated α-states, and whether we could observe centrophilic transposons forming satellite-like repeats, as reported previously^63,91–97^. Consistently, we could observe such structures derived from centromeric transposons in three distinct arrangements; (i) tandem duplication of intact transposons (**Fig. 5a-5c**), (ii) formation of satellite-like arrays from a single transposon sub-sequence (**Fig. 5d**), and (iii) tandem arrays built from mosaics of different transposon types (**Fig. 5e**).

**Figure 5.**
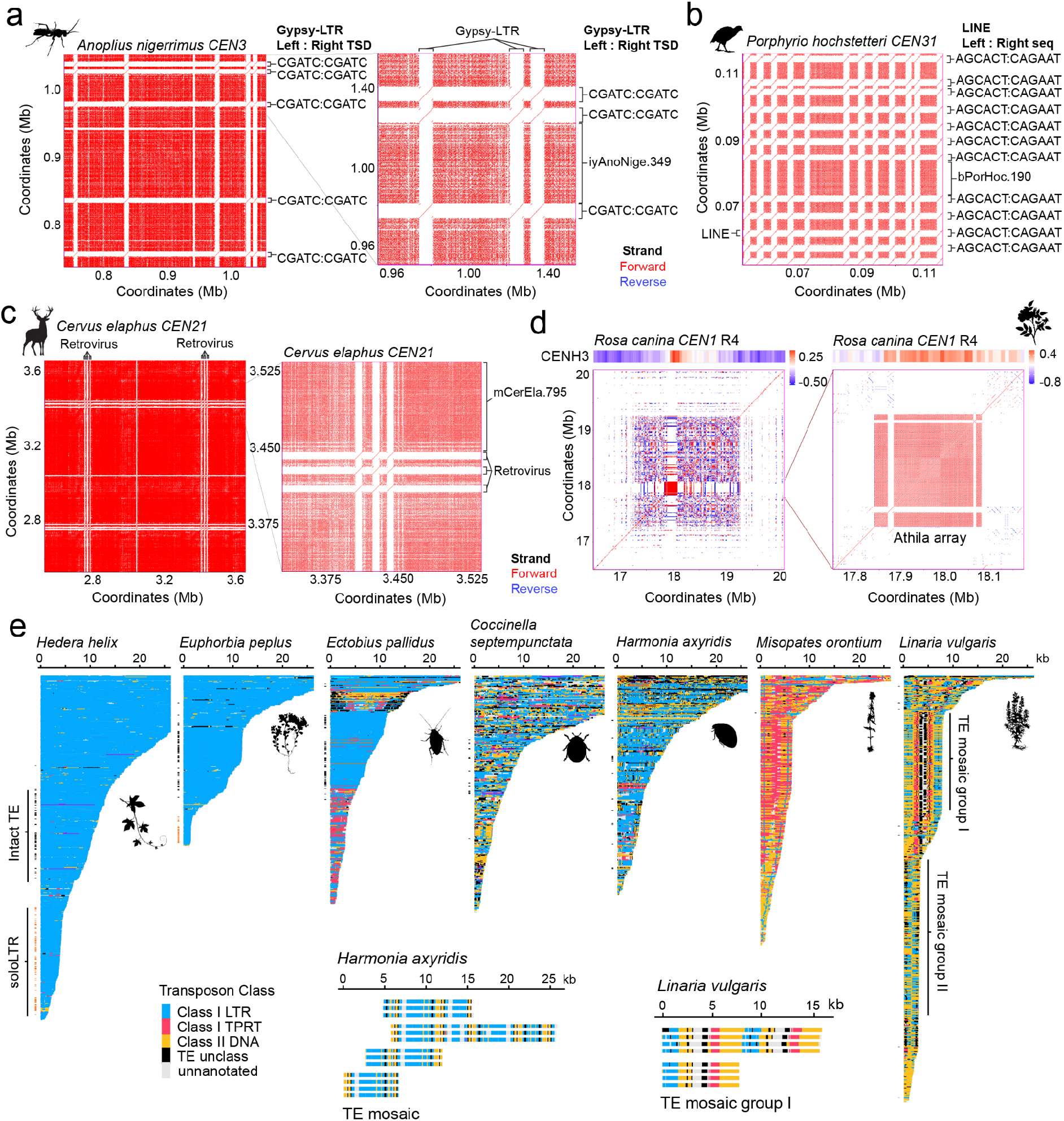
Formation of tandem repeats by centrophilic transposons. **a.** Sequence identity dot plot showing a region of centromere 3 from *Anoplius nigerrimus* with multiple copies of an LTR-Gypsy element on the same strand as the surrounding iyAnoNige.349 satellites that share identical target site duplications (TSDs). Red and blue shading indicate similarity on the forward or reverse strand, respectively (all detected homology is in plus strand). **b.** As for a, but showing a region of centromere 31 from *Porphyrio hochstetteri* (Gruiformes, Chordata), where multiple copies of a LINE retrotransposon are found within bPorHoc1.190 satellite repeats. Printed to the right are the sequences that immediately flank the LINE copies, upstream and downstream. **c.** As for a, but showing a region of *Cervus elaphus CEN21*, where a tandem cluster of three retroviruses (zoom on right) is itself duplicated on the same strand 291 kb away (visible in left panel). **d.** As for a, but showing centromere 1 from the R4 subgenome of *Rosa canina* (Rosales, Plantae), and a close-up of a 238.7 kb tandem repeat array composed of Athila sub-sequences^41^. Above each dot plot, is a heat map showing log_2_(CENH3/H3) enrichment over the same region^41^. **e.** Interruptions in the satellite arrays of indicated species are shown, ordered according to the physical length of the interruption in kilobases (capped at 25 kb), and coloured according to their transposon superfamily content. Satellite interruptions whose length matches intact elements and soloLTRs are indicated by black and orange tiles to the left of each plot. Brackets in *L. vulgaris* indicate two different groups of structured transposon-mosaic interruptions. Insets are blow-ups of satellite interruptions in *H. axyridis* and *L. vulgaris* containing transposon mosaics, including four interruptions in *L. vulgaris* that contain tandem repeats of the group I transposon mosaics.

As an example of the first state, in *Anoplius nigerrimus* five high identity Gypsy-LTR elements are located within an iyAnoNige.349 satellite array, on the same strand, which share identical target site duplications (TSDs) (**Fig. 5a**). This is consistent with a single transposon integration followed by copying via host recombination. Equivalent patterns were observed for 11 LINE elements in *Porphyrio hochstetteri*, and 6 retroviruses in *Cervus elaphus* centromere satellite arrays (**Fig. 5b-5c**). In the second class of tandem-transposon structure, sub-sequences from a single type of transposable element generate satellite-like repeats. For example, *Rosa canina* centromeres contain large tandem repeat arrays (238.7 kb on chr1, and 43.0 kb and 55.9 kb on chr4), comprised of 2.2 kb (chr1) and 1.6 kb (chr4) monomer units derived from Athila retrotransposons (**Fig. 5d**). These Athila-derived tandem arrays are strongly enriched for CENH3 binding^41^ (**Fig. 5d**), demonstrating that transposon-derived tandem repeats can be functional centromeric DNA.

Finally, we searched for tandem repeats derived from transposons when analysing interruptions of the satellite arrays. Some species, for example the plants *Hedera helix* and *Euphorbia peplus*, lack such structures and instead predominantly contain insertions of the same LTR retrotransposon superfamily, including intact elements and soloLTRs that perfectly match the length of the satellite interruption (**Fig. 5e**). In other species, such as the insects *Ectobius pallidus*, *Coccinella septempunctata*, and *Harmonia axyridis*, the satellite arrays are interrupted by more diverse superfamilies of transposons, and in addition contain small numbers of transposon-mosaics with tandem character (**Fig. 5e**, see inset for *Harmonia*). The transposon-mosaic pattern was strongest in the plants *Linaria vulgaris* and *Misopates orontium*, where we observed multiple interruptions containing shared transposon-mosaics (>200 in *L. vulgaris*), often arranged in tandem, and maintaining a consistent strand orientation that matches the surrounding satellites in each centromere (**Fig. 5e**). These observations support that transposons can be converted into satellite-like repeats by recombination at varying scales within plant and animal centromeres, providing pathways for satellite regeneration from centrophilic transposons, and an evolutionary route to connect Ω and α architectures.

### Independent origins of holocentricity with diverse underlying sequence architectures

Eleven angiosperms and 31 insects within our genome sample belong to known holocentric taxa^5,24,34,35,40^ (**Fig. 1a** and **Supplementary Table 1**). The angiosperm holocentrics are from two monocot families; Cyperaceae (n=9) and Juncaceae (n=2) in the Order Poales (**Fig. 1a** and **Supplementary Table 1**). These families include species previously observed to utilise periodic satellite repeat arrays as holocentromeres^35,40,98^. Consistently, CAP identified periodic satellite arrays in all but one of these species, with 5 monotypic, and 5 ditypic genomes (**Fig. 6a** and **Supplementary Table 2**). Within Cyperaceae, three *Schoenoplectus* species show the presence of ditypic ∼180 and ∼58 bp satellites that share 29% and 12% of 17-mers between species respectively, but whose stoichiometry varies between the genomes (**Fig. 6a**). Similarly, in the *Luzula* genus, related 125 and 104 bp satellites occur in periodic arrays, with varying stoichiometry between *L. pallescens* and *L. sylvatica* (**Fig. 6a**). For all holocentric angiosperms, we could identify one or more CENH3 orthologs (**Fig. 6a**).

**Figure 6.**
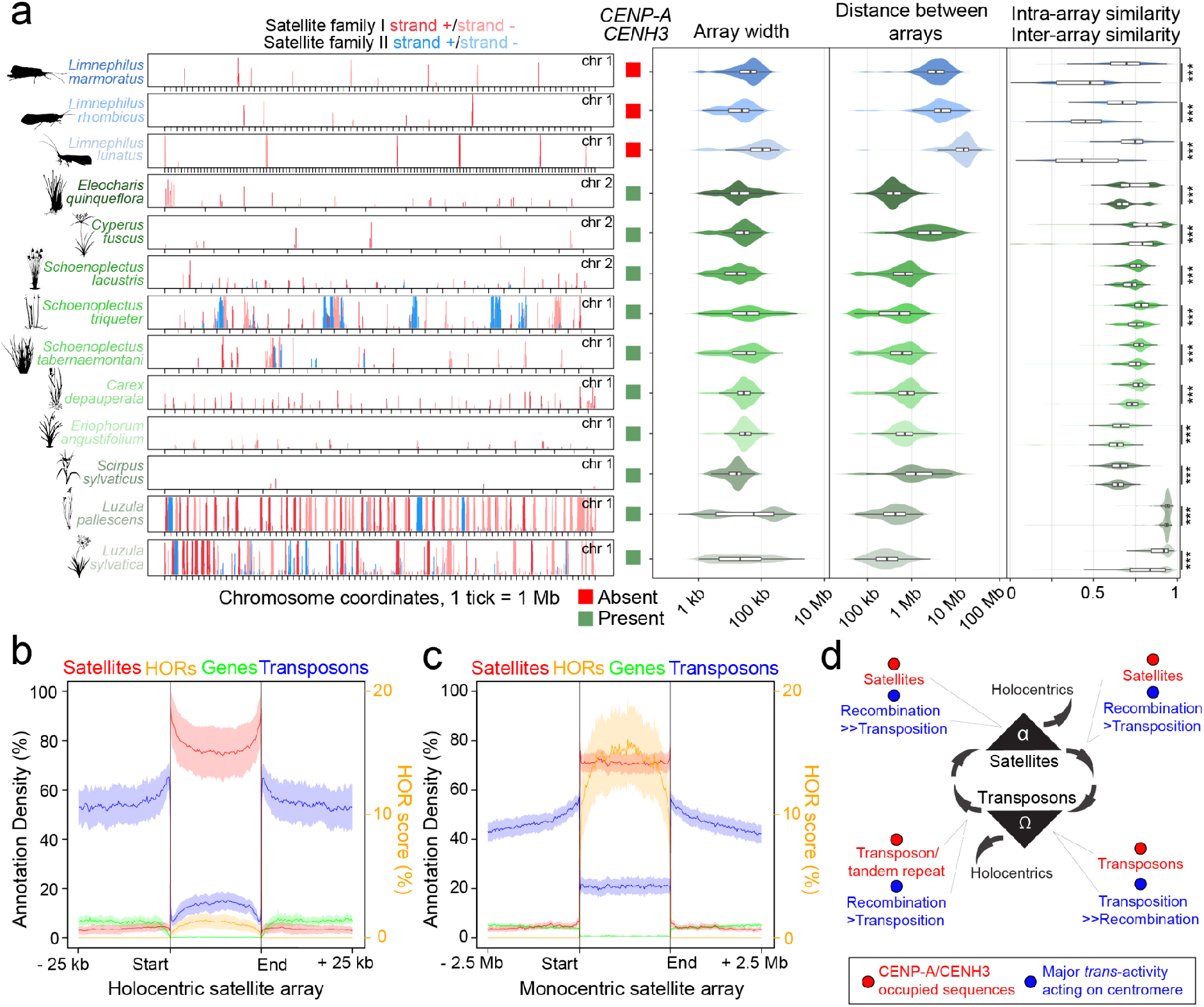
Independent origins of holocentricity with diverse underlying sequence architectures. **a.** Representative chromosomes from insect (3 *Limnephilus*, Trichoptera) and monocot (8 Cyperaceae and 2 Juncaceae) holocentric species showing periodically distributed satellite arrays, with coloured shading to distinguish array strands. Five species are ditypic, where the two satellite repeat families are shown in red and blue shading, respectively. The ticks on the x axis represent 1 megabase (Mb) distances. Shown to the right are boxes indicating the presence (green), or absence (red), of *CENH3/CENP-A* genes in the cognate genome assembly. To the right, for each assembly, are violin plots showing the width of the satellite arrays (bp), the distance between the arrays, and levels of satellite similarity within (intra, upper), and between (inter, lower) satellite arrays. For species with two satellite families, the properties were calculated independently and the results merged. **b.** Plots of scaled annotation features over holocentromeric satellite arrays for the combined species shown in a, showing satellite (red), gene (green), and transposon (blue) annotation density within scaled arrays and 25 kb upstream and downstream, and averaged HOR scores for the centromeric repeats (orange). Shading represents a 95% confidence interval for each feature. **c.** As for b, but analysing the scaled arrays of monocentric species with satellite arrays and 2.5 Mb flanking regions. **d.** A recurrent model for centromere evolution, with cycles between α-state satellite and Ω-state transposon architectural states, and independent evolution of holocentric architecture. Annotated at four stages are the identity of the CENP-A/CENH3 occupied DNA sequences shown in red, and the major *trans*-activity we consider as shaping the centromere sequences in blue; either host homologous recombination, or centrophilic transposition.

Within the insects, our sample includes holocentrics from four independent clades; Odonata (n=2), Hemiptera and Psocodea (n=6), Dermaptera (n=1), and Trichoptera and Lepidoptera (n=22) (**Fig. 1a** and **Supplementary Table 1**). Periodic or monocentric satellite repeat patterns were absent in the majority of the insect species, as observed in the lepidopteran insect *Bombyx mori*^54,99^. Periodic satellite repeats were restricted to three species within the *Limnephilus* genus (Trichoptera) (**Fig. 6a**). The mean satellite array sizes in the *Limnephilus* holocentrics are comparable to those observed in plants (iiLimRhom.187=29.5 kb; iiLimMarm.205=52.1 kb; iiLimLuna.249=134.5 kb) although, the density of the arrays per megabase was lower (0.07-0.20 arrays/Mb), than seen in the plant holocentrics (0.89-2.11 arrays/Mb) (**Fig. 6a**). Within Lepidoptera, *Pieris napi* chromosomes contain ilPieNapi.667 and ilPieNapi.79 inter-mixed ditypic arrays that are reminiscent of α-state satellite architecture; although Hi-C profiles do not indicate a monocentric organization (**Extended Data Fig. 14**). This may explain previous cytological observations of focal ‘centromeric’ constrictions in *Pieris*^100^. In contrast to the plant holocentric species, we did not identify CENP-A orthologs in the insect holocentric genomes (**Fig. 6a**), consistent with previous observations^24,34^.

For the angiosperm and *Limnephilus* genomes with periodic satellite arrays, intra-array similarity was always higher than inter-array similarity (Wilcoxon rank-sum test, Benjamini– Hochberg corrected across species; all adjusted *P*<0.001) (**Fig. 6a**), implying that recombination is largely restricted within single arrays. We quantified HORs within the holocentromeric satellite arrays and observed that HOR scores were highest in the array centres and declined towards the edges, which was reminiscent of patterns observed within larger monocentric satellite arrays (**Fig. 2h** and **6b-6c**). A further similarity between holocentromeric and monocentromeric arrays is that they show an increase in transposon density, and a decrease in gene density at their boundaries (**Fig. 6b-6c**). Together these data indicate convergent evolution of holocentricity, associated with or without periodic satellite arrays, and with or without loss of CENH3/CENP-A coding in plants and animals.

## Discussion

Centromeres are among the most rapidly evolving regions of eukaryotic chromosomes^1,2,6–9,51^. For example, we quantified a satellite repeat identity half-life decay of ∼12 Mya, and captured a centromere architectural transition in *Geum* that occurred in the last 2 million years. Despite high sequence diversity and fast evolution, centromeres form recurrent architectures typified by repetitive structures^1,2^; which we broadly characterise as α-state satellite and Ω-state transposon organisations (**Fig. 6d**). Both centromere structures create nested repeats, although α-state satellites tend to form on the same strand, reflecting mechanisms of host recombination; whereas Ω-state transposon repeats tend to occur on both strands, reflecting independent transposon integration events. Within plants and animals, we observe that α- and Ω-states are phylogenetically scattered, and we model their evolution as a recurrent cycle, with architectures recurrently emerging from one another (**Fig. 6d**). Although we endeavoured to analyse centromeres with a wide phylogenetic distribution, we have limited sampling of microbial eukaryotic lineages, which are known to contain significant centromeric novelty^26,101–103^. Future surveys should more comprehensively map centromeres in these clades.

We observe that the α-state satellites are frequent in plants and animals. Satellite monomers tend towards nucleosomal length (∼147 bp), but with a diversity of primary sequences and base compositions. Satellite higher order repeats are conserved across α-state centromeres, showing similar regimented and heterogeneous HOR patterns, higher intra than inter-chromosome satellite similarity, and spatial enrichment within the centre of the arrays. These patterns are consistent with copy number-dependent recombination processes generating the HOR structures. We observed that regimented HORs with long range periodicity, as typified in human alpha-satellite arrays^7,31,51,81^, are common across eukaryotes. The widespread occurrence of HORs is consistent with DNA breakage and homologous repair and recombination within the centromeres, which could be either mitotic or meiotic, and using either a sister chromatid or a homologous chromosome as a repair template^8,81,104^. For example, the Kinetochore Associated Recombination Machine (KARMA) hypothesis and related ideas propose that CENP-A/CENH3 chromatin is unstable, DSB-prone, and/or directly recruits recombinases, which drives rapid sequence evolution^8,81,104^. Centromeres may also be zones of DNA breakage due to mechanical forces during segregation, or replication stresses, that promote instability^105^. An alternative possibility is that nested repeat arrays form structures *in vivo* that form stronger surfaces for kinetochore attachment and therefore may reflect adapted structures for chromosome segregation.

Centromere satellite arrays show a gradation of transposon invasion. We observed a range of transposon lineages within the centromeres that use divergent integration mechanisms, supporting multiple, convergent adaptations to the centromeric niche. Tal1 provides a clear example of centrophilic invasion where the integrase C-terminus has adapted to target CENH3 chromatin^106^, and small changes to HIV-1 integrase amino acid sequence can confer centrophilic activity^107^. Selection for transposon centrophilic adaptation may occur for multiple reasons. First, centromere insertion may mitigate negative host-level selection by avoiding disruption of protein-coding genes, which are rare in the centromeres. Further, as centromeric DNA sequence is variable in terms of size and structure, these regions may represent a relatively tolerant integration environment. Second, centromeric integration may assure vertical segregation of the transposon via linkage to the centromere. In species with asymmetric meiosis (which includes female meiosis in most animals and plants), where only one of four meiotic products survives, there is also an opportunity for selfish sequences to bias their own segregation into the surviving egg cell^108–112^. Hence, centromere sequence variants, including centrophilic transposons, may cause meiotic drive to promote their own genetic transmission.

In cases where centrophilic transposons become sufficiently active, and further, the transposons themselves become the location of CENP-A/CENH3 loading, we propose the Ω state emerges. For example, Cereba-Quinta Gypsy-LTR chromoviruses in *Triticum monococcum* form centromeric clusters, and their LTR repeats are also the location of CENH3-loading^12,113^. However, if excess mutation of autonomous copies occurs, or transposon silencing becomes too effective, the requisite rate of transposon increase will not be sustained. Due to the inherent instability of centromeric chromatin proposed in the KARMA hypothesis, CENH3/CENP-A occupied transposon sequences may then undergo instability and trigger satellite formation in plants and animals (**Fig. 6d**). Supporting this, we observe satellite-like, often tandem, structures of centrophilic transposons in multiple species, consistent with previous reports^63,91–97^. We did not observe satellite architectures in fungi, where transposon centromere architectures were common. This suggests that fungi may be unable to generate or maintain satellite arrays, or that satellite arrays are selected against in this lineage.

The evolutionary forces that drive centromere architectural transitions may include centrophilic transposition, host recombination, and meiotic drive or transmission distortion. The balance of these forces may push centromeres from one architectural state to another (**Fig. 6d**). One possibility is that centromere sequences are fast-evolving due to a lack of constraint, as centromere identity is fundamentally epigenetic. However, an alternate possibility is that while centromere primary sequences are not functionally constrained, repeat architecture is conserved across plants and animals, and nested repeat structures may form more effective kinetochore surfaces that can selfishly compete during chromosome segregation. We further speculate that KARMA-mediated recombination regimes may represent, in part, a genomic defence against driving sequences. Continual rewriting of centromere sequences by homologous recombination could erase or equalise driving sequences that attempt to exploit centromere function. The convergent evolution of dispersed holocentromeric centromere function could itself further represent a defence against drive. For example, by distributing spindle attachment across many loci, holocentric chromosomes may avoid the single centromeric target that selfish driving elements aim to exploit. Further pangenomic exploration of species diversity across eukaryotes will provide new insights into centromere evolutionary dynamics, and the selective drivers that shape their evolution.

---

**Extended Data Figure 1.**
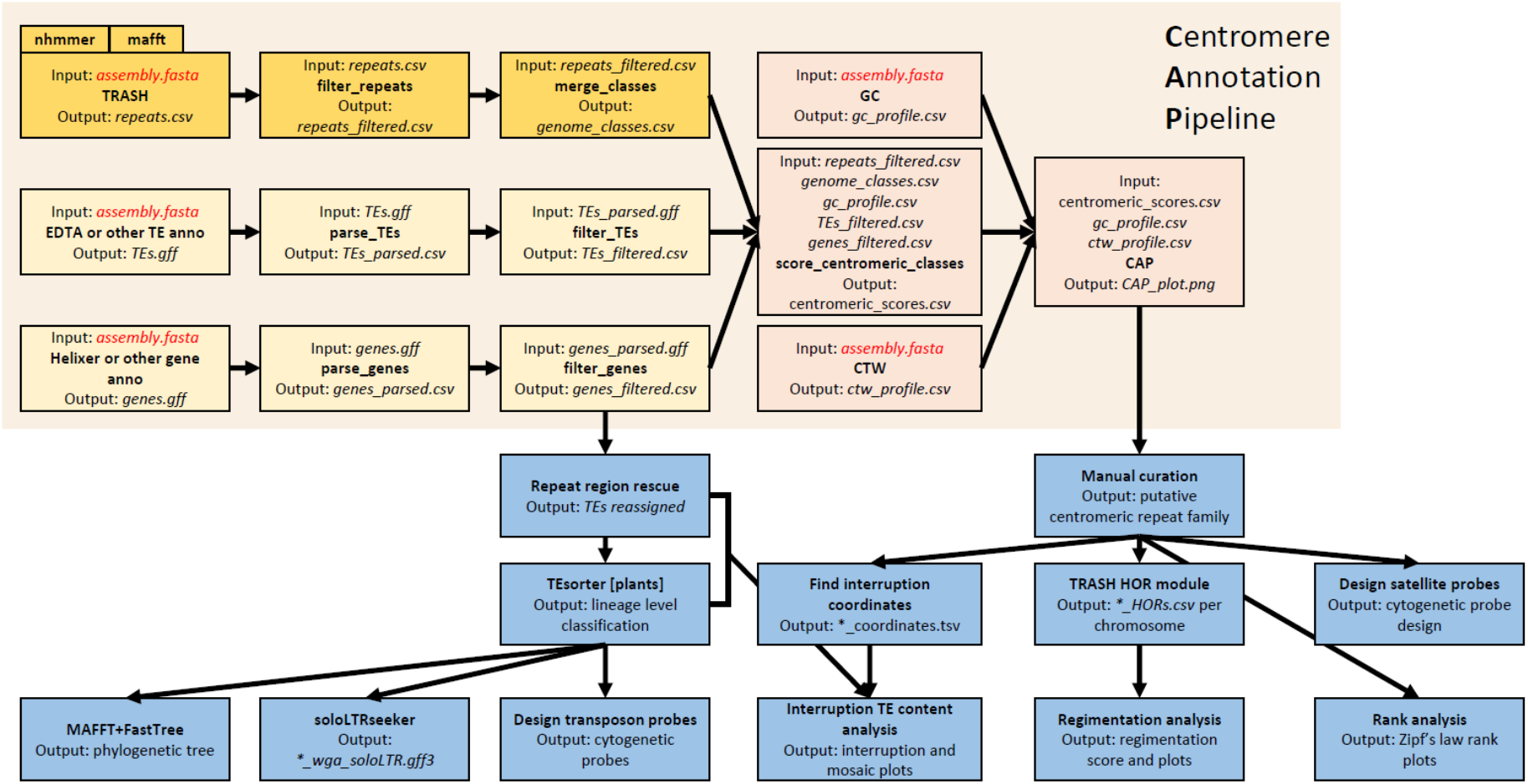
Centromere Annotation Pipeline (CAP). A diagrammatic overview of the Centromere Annotation Pipeline (CAP) and downstream workflows. CAP (salmon boxes) receives tandem repeat annotation from TRASH^46^ (orange), as well as gene and transposon annotation (yellow boxes). The CAP annotation, together with available Hi-C data, literature surveys, and manual inspection were used to assign a centromere architecture classification to each genome. The blue boxes represent the main steps of downstream analyses, following CAP. The scripts associated with the steps in this diagram can be found in the project GitHub repository (https://github.com/vlothec/eukaryotic_centromere_architecture).

**Extended Data Figure 2.**
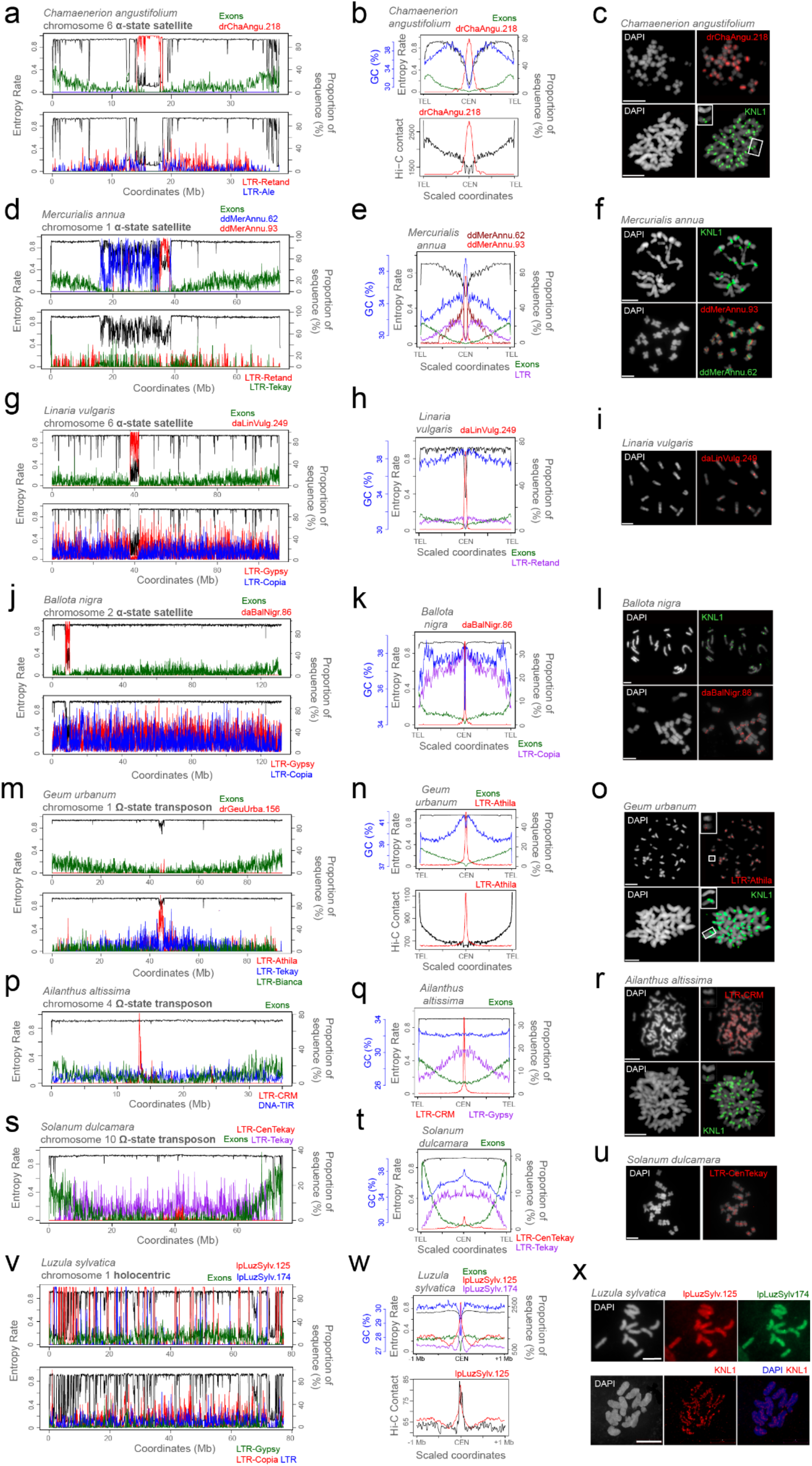
FISH and immunocytological validation of CAP centromere predictions. a-l show validation for α-state satellite CAP classified genomes, m-u show validation for Ω-state transposon classified genomes, and v-x shows validation for a holocentric genome. **a.** Chromosome 6 of *Chamaenerion angustifolium* is plotted showing entropy rate (black), and the proportion of sequence that was drChaAngu.218 satellite (red), exon (green), and Retand (red), and Ale (blue) LTR retrotransposons. **b.** Data as in a for *C. angustifolium*, but analysed in scaled windows between the centre of satellite arrays and the telomeres, with %GC (blue), drChaAngu.218 satellite (red), exons (green), and entropy rate (black). Beneath is a plot of Hi-C intra-chromosome contact frequency analysed in the same windows, together with drChaAngu.218 density. Hi-C contact frequency was extracted in 50 kb bins. **c.** Cytogenetic analysis of *C. angustifolium*, with DNA stained by DAPI, immunostaining for KNL1 (green), and FISH for the drChaAngu.218 satellite repeat (red). Scale bars=5 µM. **d.** Chromosome 1 of *Mercurialis annua* is plotted showing entropy rate (black), and proportion of sequence that was ddMerAnnua.62 (blue) and ddMerAnnua.93 satellite (red), exon (green), and Retand (red) and Tekay (green) LTR retrotransposons. **e.** Data as in d for *M. annua*, are analysed in scaled windows between the centre of ddMerAnnua.93 arrays and the telomeres with %GC (blue), ddMerAnnua.62 (dark red) and ddMerAnnua.93 satellite (red), exons (green), LTR transposons (purple), and entropy rate (black). **f.** Cytogenetic analysis of *M. annua* is shown as in a with DNA stained by DAPI, and FISH for the ddMerAnnua.62 (green) and ddMerAnnua.93 (red) satellite repeats and KNL1 immunostaining (green). **g.** Chromosome 6 of *Linaria vulgaris* showing entropy rate (black), and the proportion of sequence that was daLinVulg.249 satellite (red), exon (green), and LTR-Gypsy (red), and LTR-Copia (blue) LTR retrotransposons. **h.** Data as in g for *L. vulgaris*, but analysed in scaled windows between the centre of satellite arrays and the telomeres, with %GC (blue), daLinVulg.249 satellite (red), exons (green), LTR-retand (purple) and entropy rate (black) shown. **i.** Cytogenetic analysis of *L. vulgaris*, with DNA stained by DAPI, and FISH for the daLinVulg.249 satellite repeat (red). Scale bars=5 µM. **j.** Chromosome 2 of *Ballota nigra* (Lamiales) is plotted showing entropy rate (black), and the proportion of sequence that was daBalNigr.86 satellite (red), exon (green), and LTR-Gypsy (red), and LTR-Copia (blue) LTR retrotransposons. **k.** Data as in j for *B. nigra*, but analysed in scaled windows between the centre of satellite arrays and the telomeres, with %GC (blue), daBalNigr.86 satellite (red), exons (green), LTR-Copia (purple) and entropy rate (black) shown. **l.** Cytogenetic analysis of *B. nigra*, with DNA stained by DAPI, immunostaining for KNL1 (green), and FISH for the daBalNigr.86 satellite repeat (red). Scale bars=5 µM. **m.** Chromosome 1 of *Geum urbanum* is plotted showing entropy rate (black), and the proportion of sequence that was drGeuUrba.159 satellite (red), exon (green), and LTR-Athila (red), LTR-Bianca (green) and LTR-Tekay (blue) LTR retrotransposons. **n.** As for b, but analysing *G. urbanum* centred on Athila arrays. **o.** Cytogenetic analysis of *G. urbanum*, with DNA stained by DAPI, immunostaining for KNL1 (green), and FISH for the Athila transposons (red) (reproduced in Fig. 4a). **p.** Chromosome 4 of *Ailanthus altissima* is plotted showing entropy rate (black), exon density (green), Gypsy LTR-CRM (red) and DNA-TIR (blue) transposon density. **q.** As in e, but for *A. altissima* analysed in scaled windows between the centre of LTR-CRM arrays and the telomeres, and showing LTR-Gypsy (purple), gene exons (green) and GC% content (blue). **r.** Cytogenetic analysis of *A. altissima*, with DNA stained by DAPI, immunostaining for KNL1, and FISH for the centromeric LTR-CRM transposons. Scale bars=5 µM. **s.** Chromosome 12 of *Solanum dulcamara*, but where purple indicates LTR-Tekay elements, and red indicates LTR-CenTekay sequences that match the FISH probe used in u. **t.** As for e, but centred on *S. dulcamara* CenTekay arrays. **u.** As for i, but analysing *S. dulcamara* with CenTekay FISH oligo probes. **v.** As for a, but showing *Luzula sylvatica* chromosome 1 as a representative of holocentric centromere architecture plotting entropy rate (black), gene exons (green), lpLuzSylv.125 (red) and lpLuzSylv.174 (blue) **w.** As for b, but analysing scaled centromere to telomere windows, we analysed 1 Mb upstream and downstream of all holocentromeric arrays. Hi-C contact frequency was extracted in 10 kb bins. **x.** As for u, but FISH in *Luzula sylvatica* for the lpLuzSylv.125 and lpLuzSylv.174 ditypic satellite repeats.

**Extended Data Figure 3.**
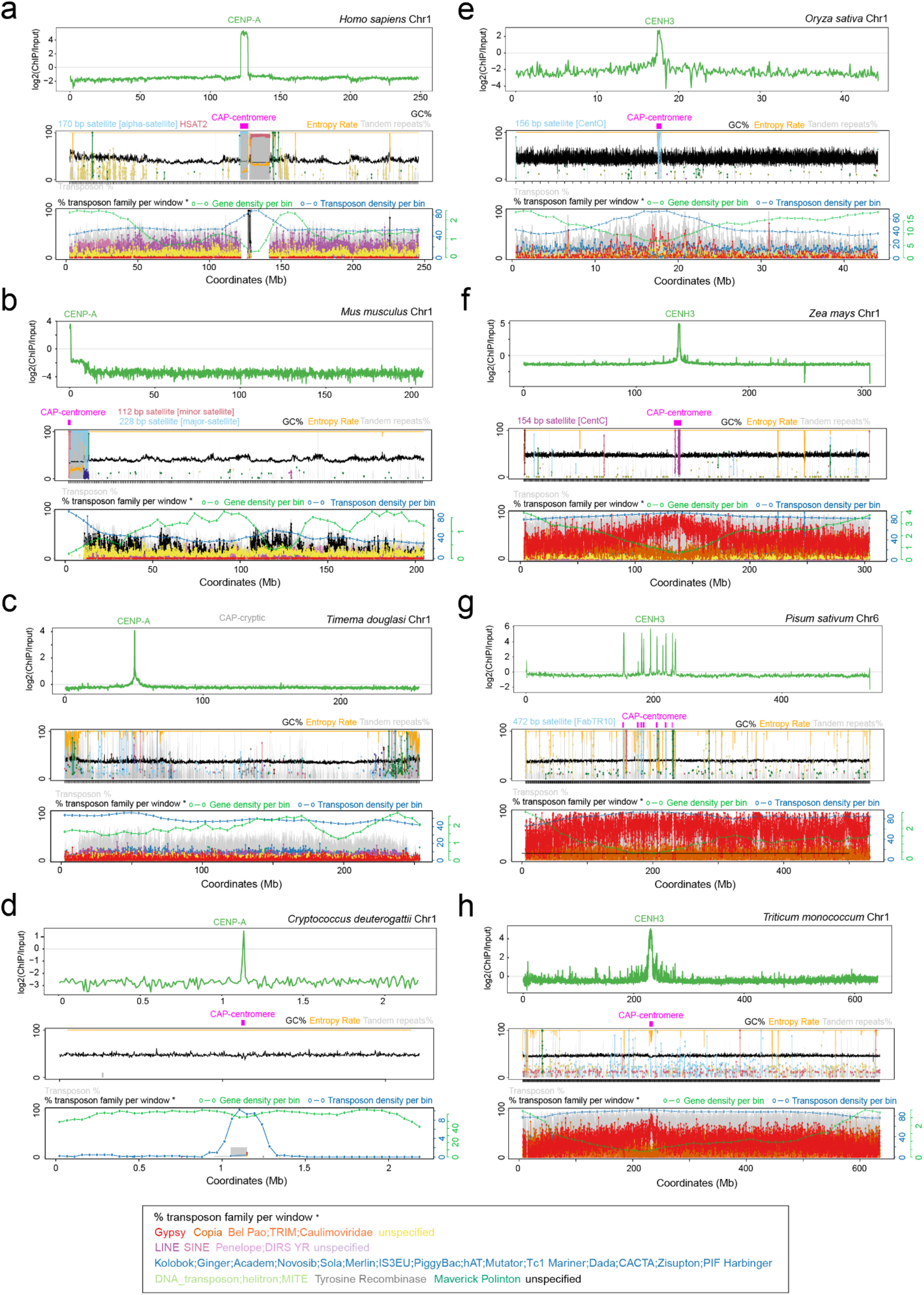
Validation of CAP centromere predictions in CENP-A/CENH3 characterised genomes. CAP annotation from representative chromosomes of 8 animal, plant, and fungal genomes, comparing centromere architecture classification with CENP-A/CENH3 ChIP-seq enrichment. For each genome, the upper panel shows log2(ChIP/input) (green) enrichment of CENP-A/CENH3. The middle and lower panels show CAP output plots that quantify tandem repeat, gene, and transposon annotations. The middle plot includes labels above (pink) highlighting regions identified as candidate centromeres by CAP. The following genomes were analysed: **a.** *Homo sapiens* CENP-A ChIP-seq (SRX255043) and input (SRX255044) data were aligned against the CHM13 (GCF_009914755.1) assembly^31^. **b.** *Mus musculus* CENP-A ChIP-seq (SRX20577194) and input (SRX20577193) data were aligned against the GCA_000001635.9 assembly^58^**. c.** *Timema douglasi* CENP-A ChIP-seq (SRX28824591) and input (SRX28824592) data were aligned against the GCA_040436065.2 assembly^61^. **d.** *Cryptococcus deuterogattii* R265 CENP-A ChIP-seq (SRX3049403) and input (SRX3049404) data were aligned against the GCA_002954075.1 assembly^33^. **e.** *Oryza sativa* CENH3 ChIP-seq (SRX27798377) and input (SRX27798382) data were aligned against the GCA_001623365.2 assembly^59^. **f.** *Zea mays* CENH3 ChIP-seq (SRX17512265) and input (SRX17512267) data were aligned against the GCA_022117705.1 assembly^60^. **g.** *Pisum sativum* CENH3 ChIP-seq (ERR9981080) and input (ERR9981081) data were aligned against the GCA_977071245.1 assembly^114^. **h.** *Triticum monococcum* CENH3 ChIP-seq (DRX799804) and input (DRX799805) data were aligned against the GCA_057380995.1 assembly^12^.

**Extended Data Figure 4.**
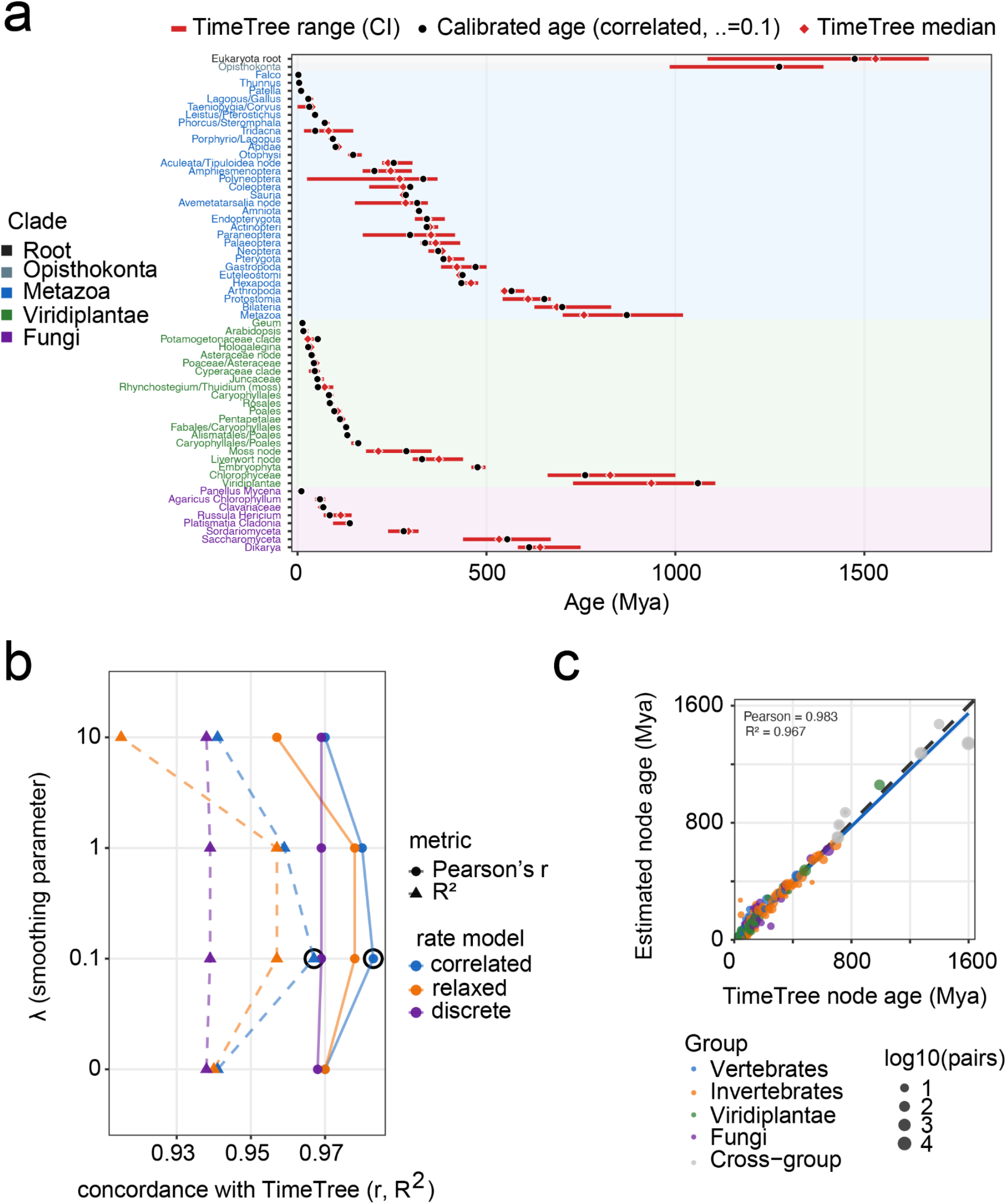
Time calibration of the species tree. **a.** Calibration constraint intervals for all 62 nodes used to time-calibrate the 325 species phylogeny. Red bars indicate TimeTree constraint ranges (min–max bounds derived from TimeTree^67^), and the TimeTree median ±20% for nodes lacking a reported range (4 of the 62 nodes). Filled circles show estimated node ages with the best tree, and diamonds show TimeTree median ages. The black circles indicate the calibrated age. **b.** Selection of the dating model. Node-age concordance with TimeTree for chronos under three rate models (correlated, relaxed, and discrete) across a range of the smoothing parameter (λ=0, 0.1, 1, and 10). Concordance is summarised by Pearson’s *r* and *R²*, each computed across the 213 node-age comparisons. The correlated-rates model at λ=0.1 gave the highest concordance (best values are circled in black). **c.** Node-age concordance with TimeTree for the best model (correlated, λ=0.1). Each point is the age of one most-recent-common-ancestor shared by the 210 species common to both trees of 325 tips, comparing our estimate (y) with TimeTree (x). The 21,945 species pairs resolve to 213 nodes, and dot size scales with the number of pairs per node (log₁₀). The dashed grey line shows 1:1 concordance, and the blue line is the linear fit.

**Extended Data Figure 5.**
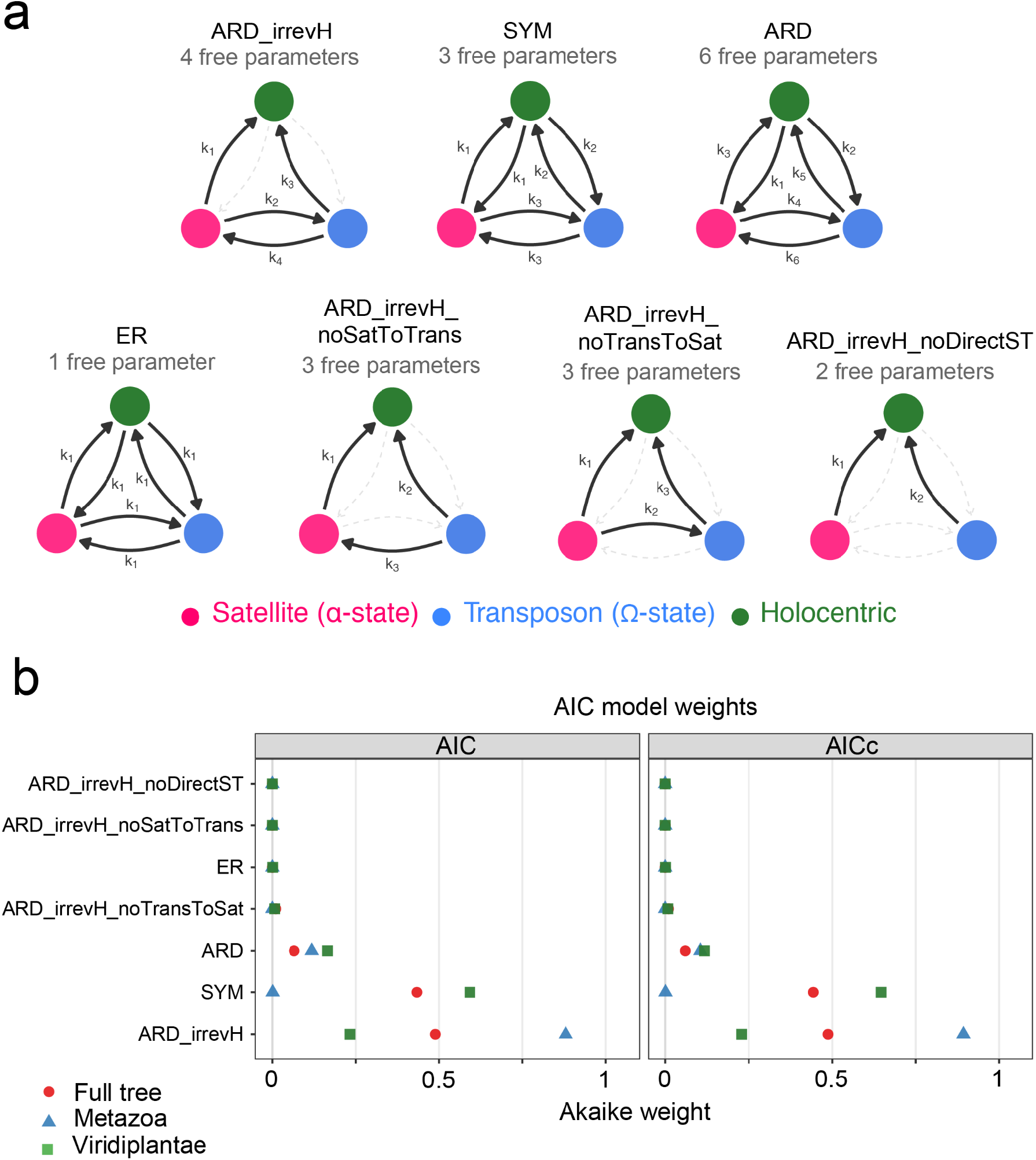
Modelling centromere evolution using ancestral state reconstruction. **a.** Transition diagrams for the Mk models evaluated in the three-state sensitivity analysis, considering holocentric (green), α-state satellite (pink), and Ω-state transposon (blue) centromere architectures. Mixed α-state/Ω-state and cryptic taxa were excluded due to low numbers of these classes. Standard models (ER, SYM, ARD) are shown, alongside three biologically motivated custom models that imposed constraints on holocentric transition irreversibility, and satellite↔transposon architecture transitions. The number of free parameters is shown below each model name. The solid black arrows indicate allowed transitions in the model (labelled ki). The dashed grey lines indicate rates fixed to zero. **b.** Akaike information criterion (AIC) and sample-size-corrected AICc weights for each model, shown separately for the full tree (red), Metazoa (blue), and Viridiplantae (green) datasets. Weights sum to 1 within each dataset and criterion.

**Extended Data Figure 6.**
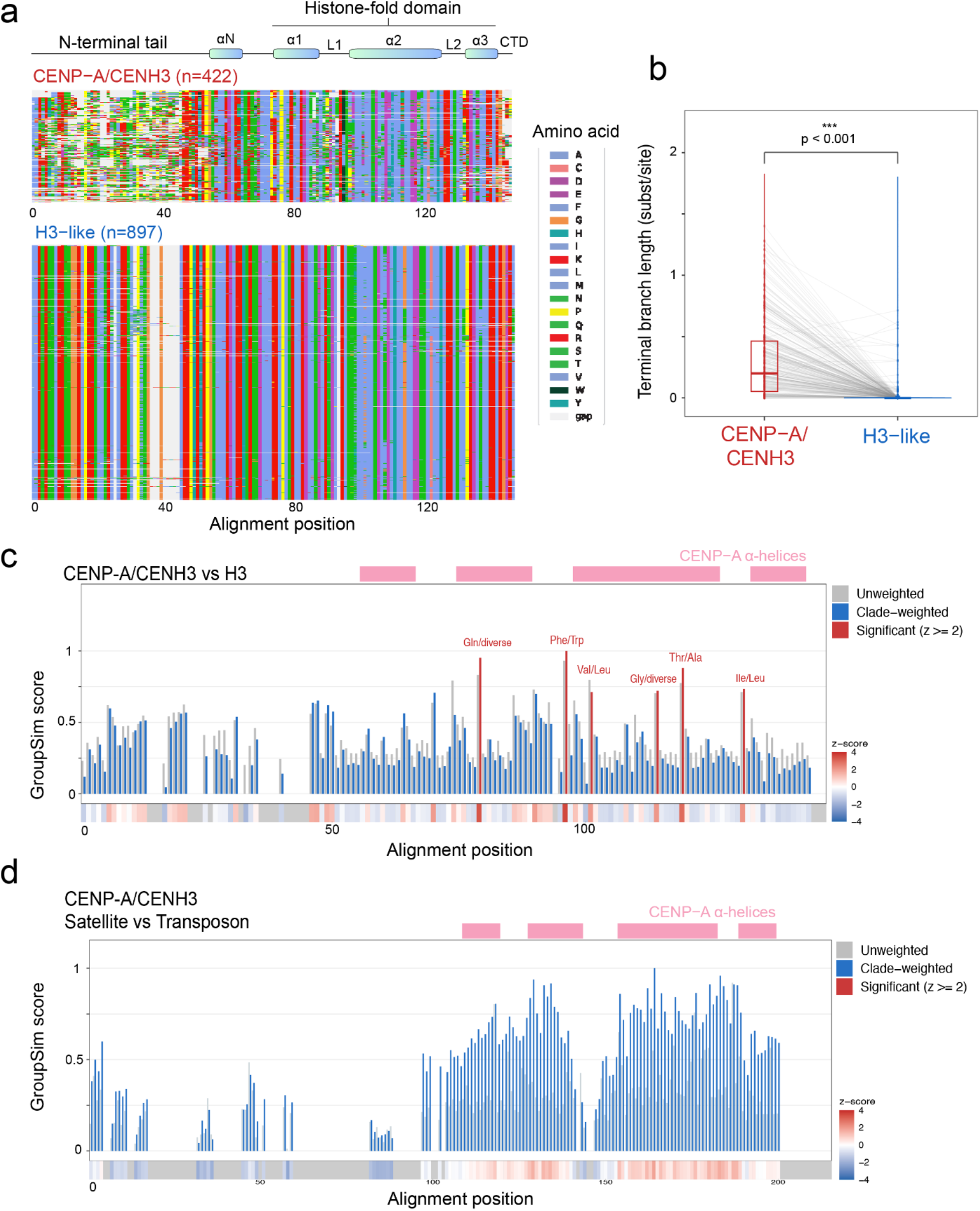
CENP-A/CENH3 protein level variation across 325 eukaryotic species. **a.** A multiple sequence alignment of CENP-A/CENH3 (n=422, upper) and H3-like (n=897, lower) sequences, visualised as an amino acid matrix, after trimming columns with >85% gaps. Amino acids are coloured according to the key shown to the right. The bar at the top indicates the predicted α-helical regions of CENP-A/CENH3, derived from AlphaFold2 structural models of *Arabidopsis thaliana* CENH3 (UniProt Q8RVQ9) using STRIDE^78^, followed by projection onto the trimmed alignment coordinates. **b.** Terminal branch distances in the CENP-A/CENH3 and H3 gene tree for paired CENP-A and H3-like sequences from the same species (n=262). Lines connect the median distances per species. CENP-A shows significantly greater RTD than H3 (Wilcoxon signed-rank test P<0.001). **c.** GroupSim^75^ specificity scores for CENP-A/CENH3 versus H3-like sequences. Grey bars indicate the unweighted mean values, whereas blue bars show clade-weighted values in which each sequence is weighted by 1/n within its broad taxonomic group to correct for uneven phylogenetic sampling. Six positions reach z≥2 (labelled in red) under clade weighting. The colour strip below shows a heat map of GroupSim z-score per position. Regions with >85% gaps are shown as grey positions. The highlighted positions in red include established sequence differences between CENP-A/CENH3 and H3^76,77^. **d.** As for c, but showing GroupSim specificity scores for CENP-A/CENH3 sequences from genomes with satellite versus transposon-based centromere architectures. No position reached z≥2 under either scheme.

**Extended Data Figure 7.**
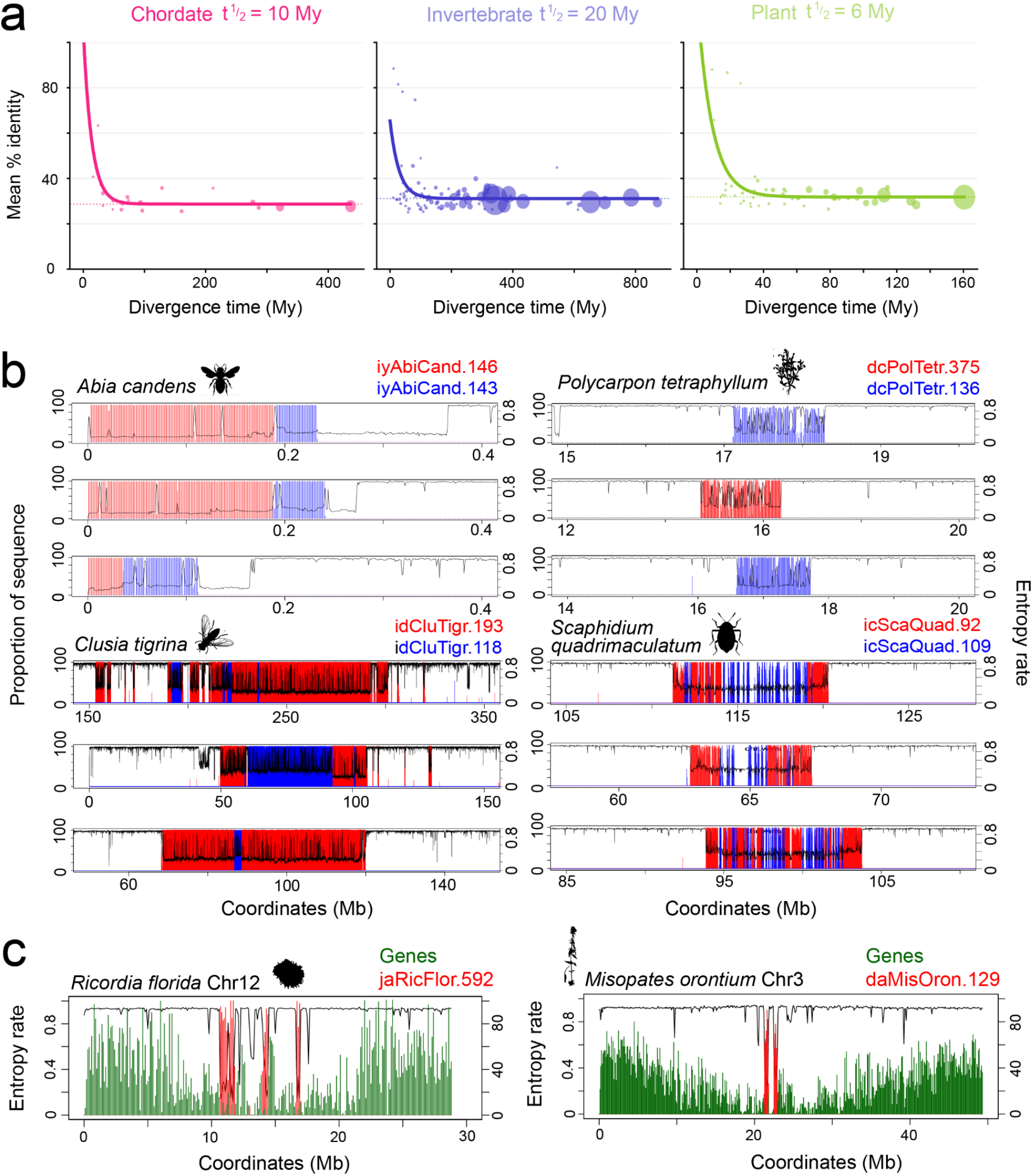
Satellite similarity and candidate metapolycentric architectures. **a.** Pairwise satellite sequence similarity as a function of species divergence time, computed by BLASTn across all satellite families within chordates, invertebrates, and plants. Each point represents the mean BLASTn-derived pairwise sequence identity between satellite families at a given node-averaged divergence time. The point size is proportional to the square root of the number of species pairs contributing. The solid lines show exponential decay fits (sim = A·exp(−λt)+C) with an empirically estimated background floor C (dotted horizontal line). Half-lives (t½ = ln(2)/λ) are indicated per clade. **b.** Representative centromere regions from species with ditypic satellites, with the position and proportion of the two families plotted in red and blue, as well as sequence entropy rate (black). **c.** Gene (green) and satellite repeat (red) density along Ricordia florida chromosome 12, and Misopates orontium chromosome 3, which show clustering of discrete satellite arrays, resembling metapolycentric architectures^39,84^.

**Extended Data Figure 8.**
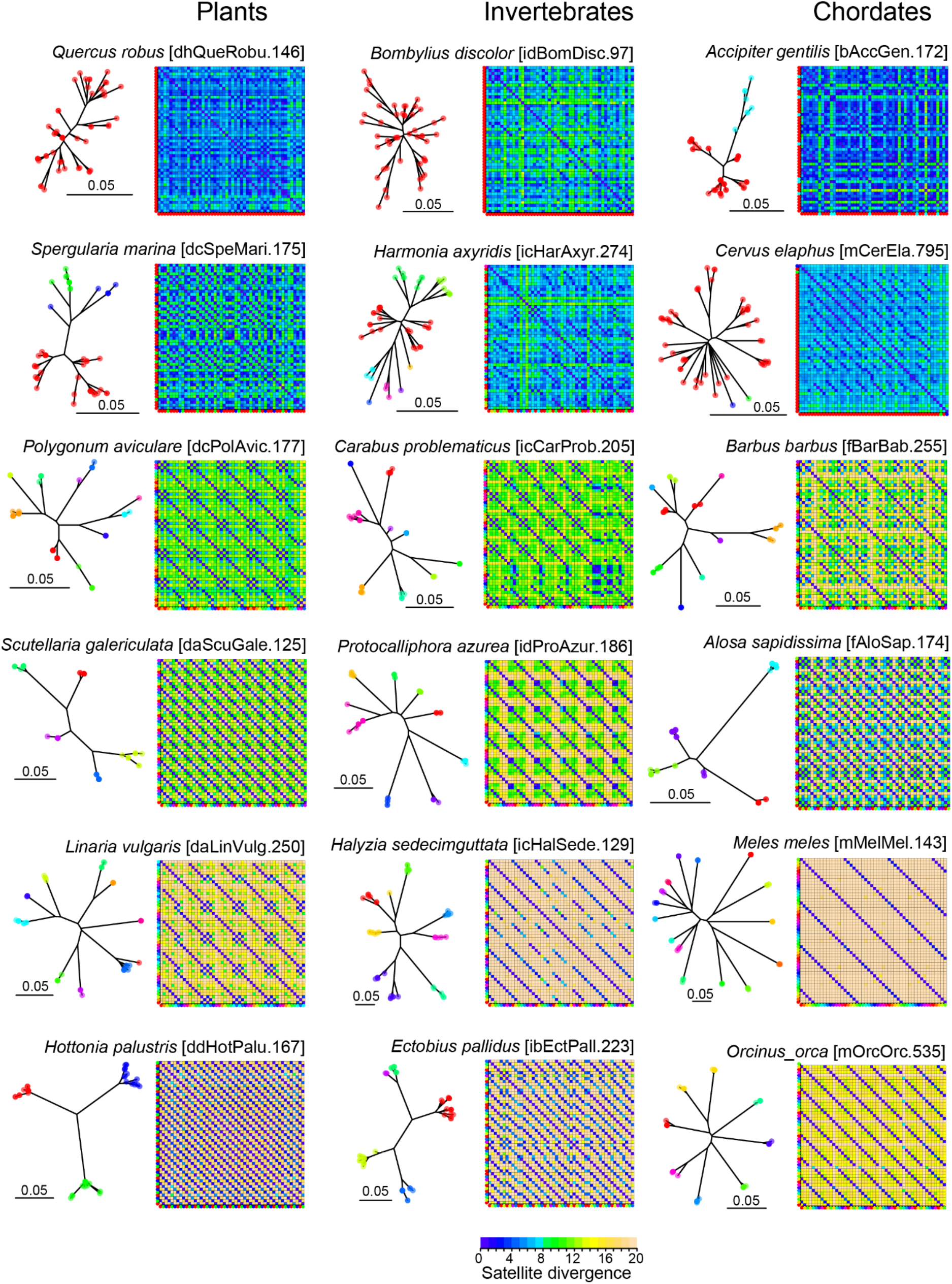
Regimented and heterogeneous satellite higher order repeats in plants and animals. For each species, a sample of 50 contiguous satellite repeats were analysed from a single centromere, and the pairwise divergence of all monomers plotted in a 2D matrix. A phylogenetic tree of the same repeats is shown to the left, where clade tips are coloured. The same clade colours are displayed along the x and y axis of the divergence 2D matrix. Each column of plots were selected from plant, invertebrate, and chordate taxa, and were chosen to represent a range of heterogeneous and regimented satellite higher order repeat organisation.

**Extended Data Figure 9.**
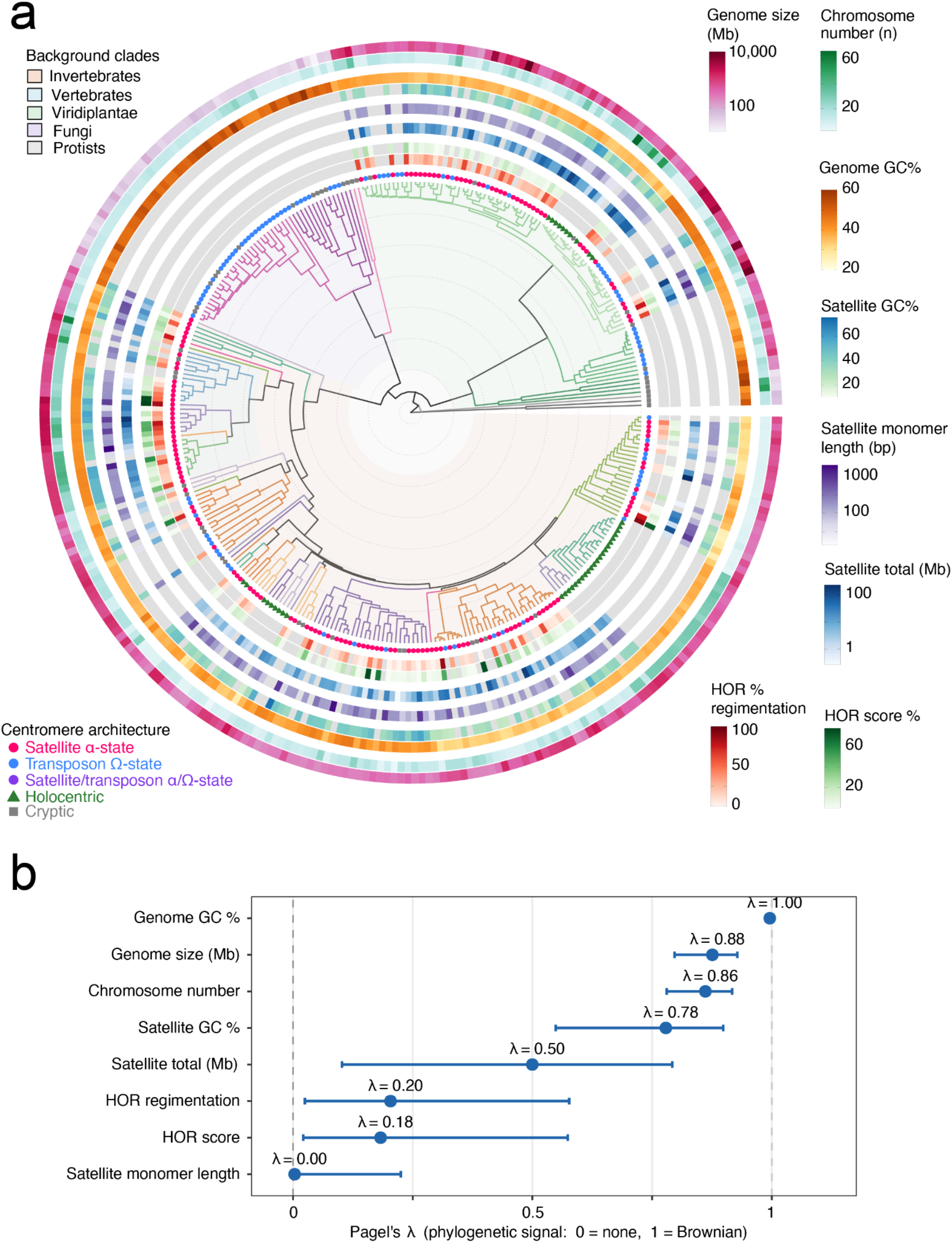
Satellite and genome properties across the eukaryotic species tree. **a.** Phylogenetic species tree generated using a BUSCO gene multiple sequence alignment using FastSpeciesTree^64^, as in Figure 1. Branch lengths are shown in millions of years (Mya) from TimeTree^67^ fossil calibration constraints, with concentric dotted rings marking 200-million-year intervals. Background tree shading colours denote major clades. CAP-derived centromere architecture classifications are indicated on branch tips as satellite α-state (pink), transposon Ω-state (blue), mixed α-/Ω-states (purple), holocentric (green), or cryptic (grey). The outer rings, from the tree outward, show heatmap quantification of: (1) HOR regimentation score (%), (2) HOR score (%), (3) satellite repeat total amount (Mb), (4) mean satellite monomer length (mean bp per family), (5) satellite GC% (6) genome GC%, (7) chromosome number, and (8) genome size (Mb). For ditypic and polytypic satellite species (those carrying two or more distinct centromeric satellite families) satellite- and HOR-derived values are taken from the single most abundant family (highest monomer copy number). **b.** Phylogenetic signal (Pagel’s λ) of genome and centromere metrics across the 325 species tree. Points represent maximum-likelihood estimates of Pagel’s λ and horizontal bars show the 95% confidence from the likelihood profile. The dashed lines mark λ=0 (no signal, showing that a metric varies independent of phylogeny), and λ=1 (the metric distribution across the tree matches the Brownian-motion expectation showing strong phylogenetic signal).

**Extended Data Figure 10.**
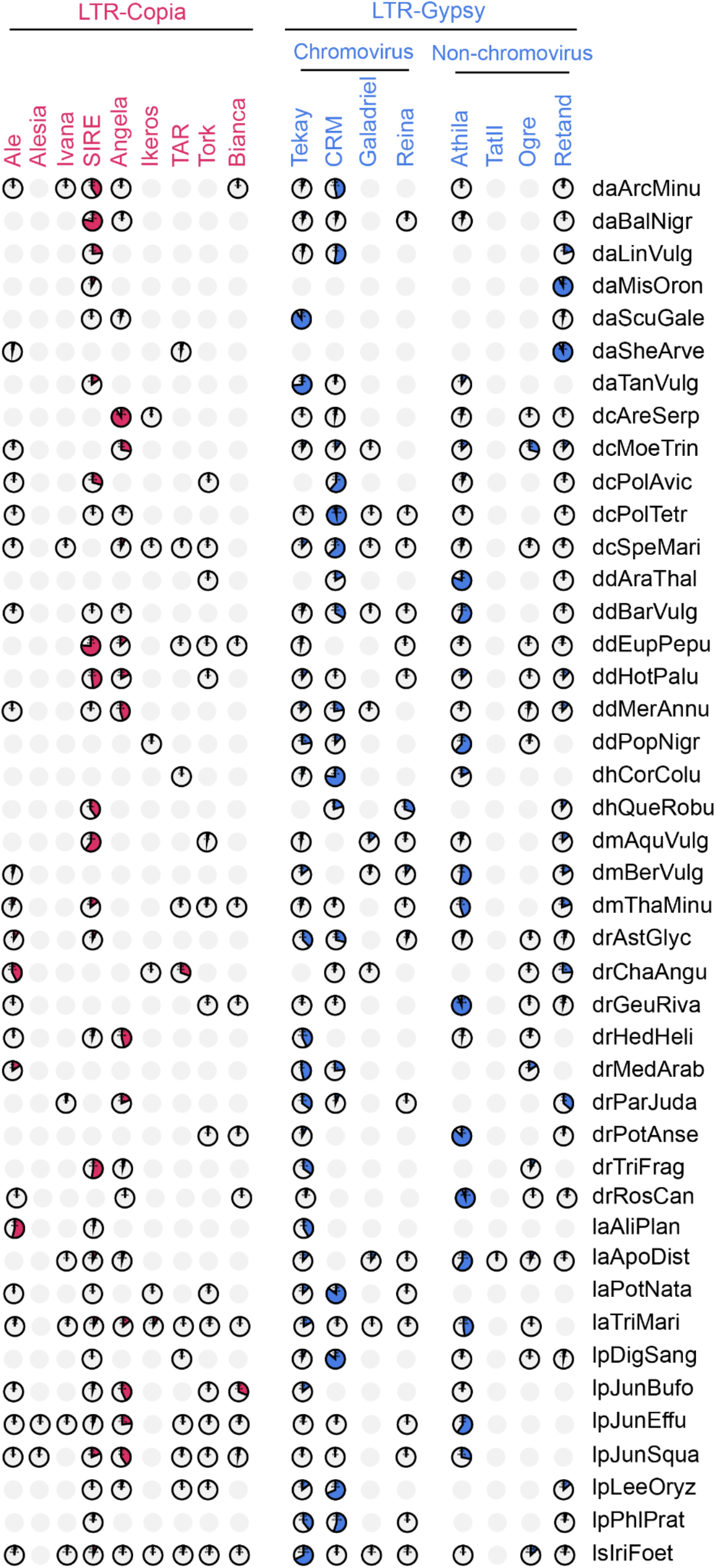
Diversity of centrophilic LTR retrotransposon lineages found in angiosperm satellite arrays. Pie charts indicating the relative proportion of Copia (red) and Gypsy (blue) LTR retrotransposon lineages found within the centromeric satellite arrays of angiosperms. The relative proportion was calculated by dividing the DNA space of every lineage within arrays, by the total DNA space of all lineages within the centromere arrays combined. Empty charts indicate the absence of an LTR lineage within the satellite arrays from that genome. The number of annotated elements (intact and fragmented) and their total length (kb) for a lineage is printed inside each pie chart. The host genome TolID is printed alongside the pie chart rows.

**Extended Data Figure 11.**
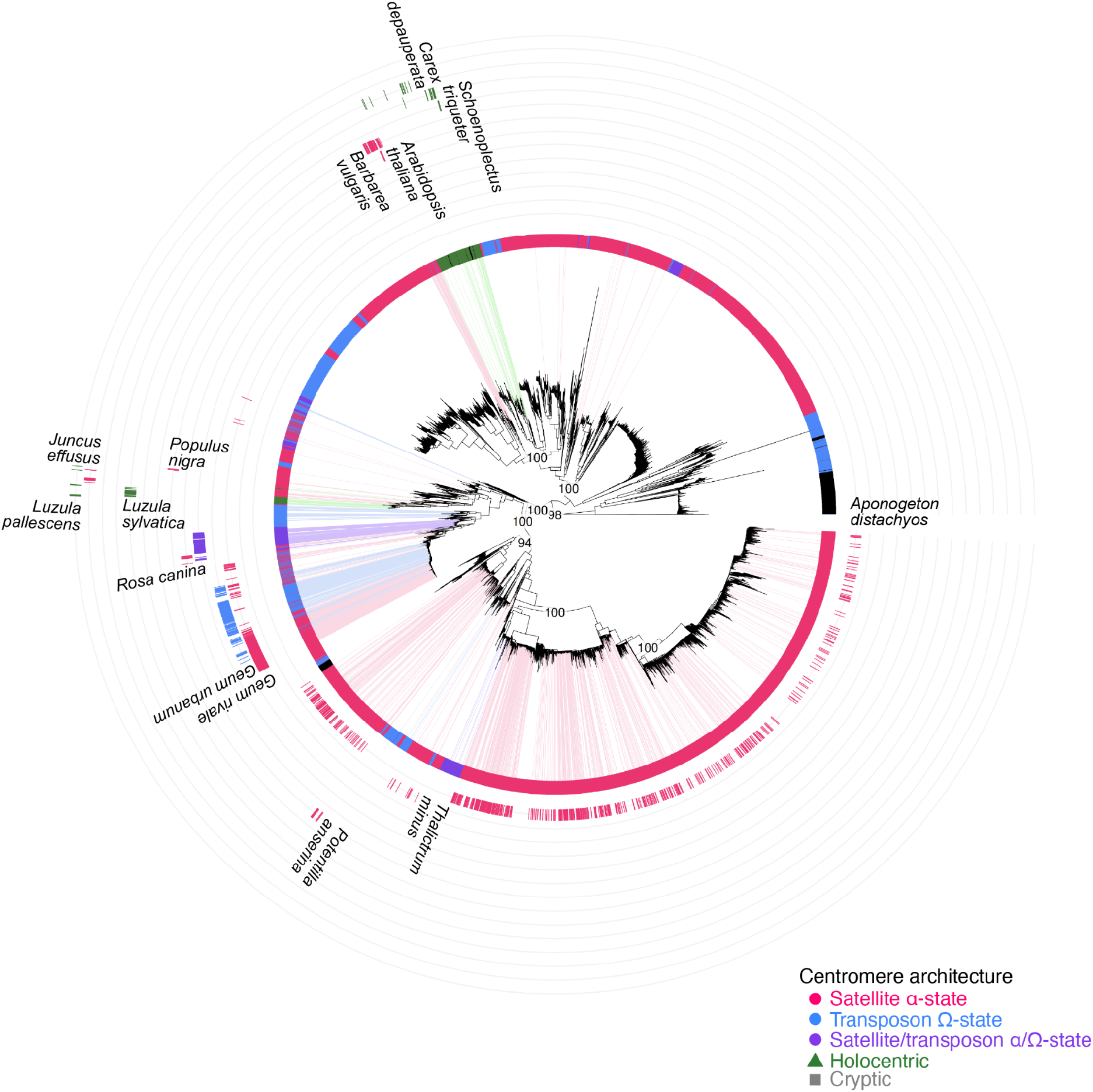
Phylogenetic analysis of Gypsy-Athila LTR retrotransposons in angiosperms. The phylogenetic tree was generated based on the alignment of five concatenated functional domains (defined by HMM profiles) of the gag, protease, reverse transcriptase, RNAaseH and integrase genes of 48,339 intact Athila elements identified genome-wide in 90 angiosperm species. Centromere architecture classification is indicated by the inner coloured ring for each species (pink for satellite α-state, blue for transposon Ω-state, purple for mixed α-/Ω-state, green for holocentric, and grey for cryptic). 2,007 Athila elements located within the centromeric satellite arrays of 13 species are highlighted in the outer rings and also with coloured shading that follows their centromere architecture classification. The outer rings also show the species TolID, to indicate independent colonizations of Athila across angiosperms. Bootstrap support of key nodes is included.

**Extended Data Figure 12.**
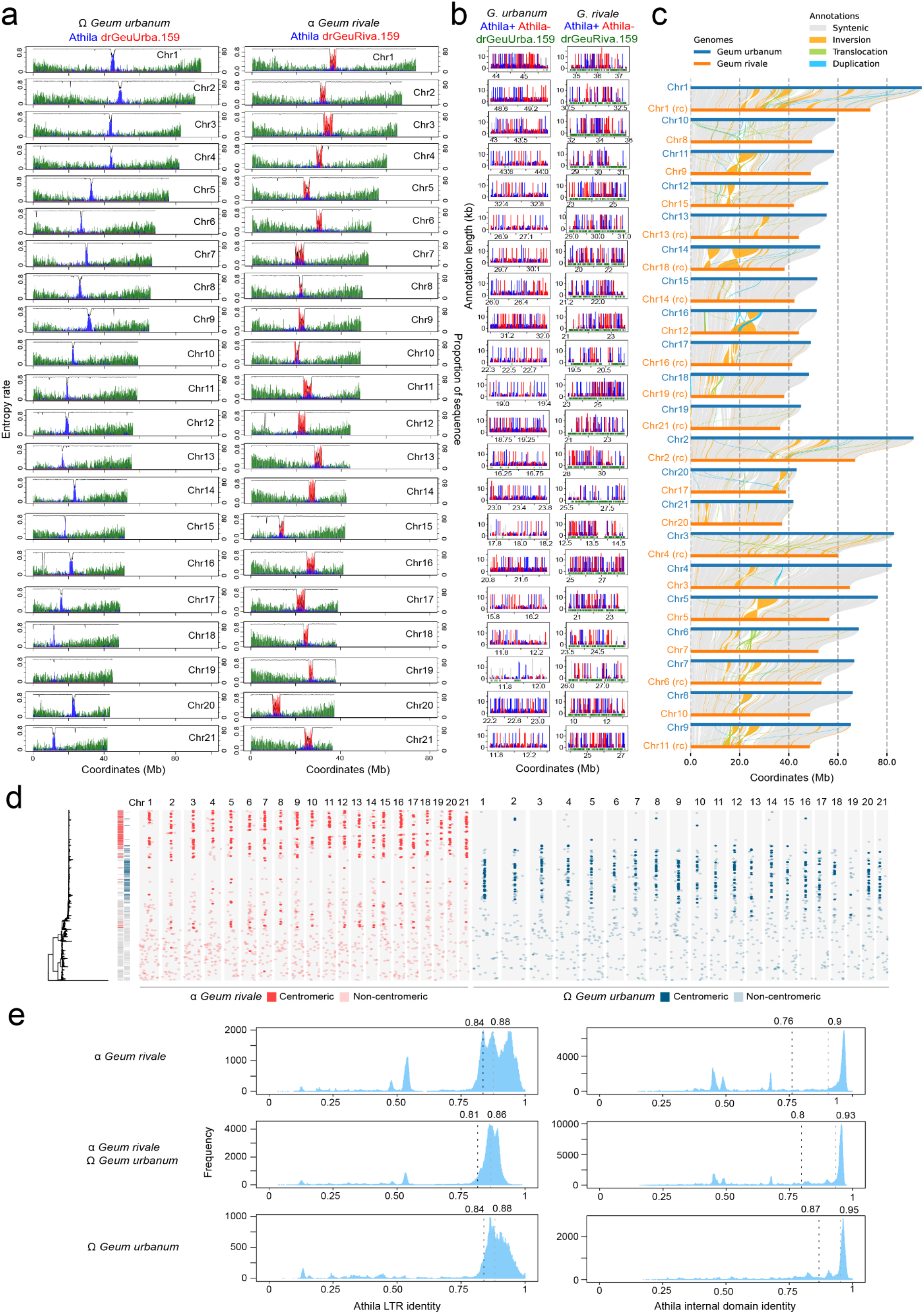
Comparison of the *Geum urbanum* and *Geum rivale* genomes and centromere architectures. **a.** Plots of all *G. urbanum* and *G. rivale* chromosomes showing the proportion of sequence corresponding to genes (green), Athila retrotransposons (blue), and 159 base pair satellite repeats (red), as well as sequence entropy rate (black). **b.** Plots of *G. urbanum* and *G. rivale* centromeres showing the position and length of Athila annotations on forward (red) and reverse (blue) strands. The position of drGeuRiva.159 and drGeuUrba.159 repeats is also indicated along the x-axis (green). **c.** *G. urbanum* and *G. rivale* chromosomes are connected by SyRI synteny diagrams, showing regions of synteny (grey), inversions (orange) and translocations (green). **d.** A phylogenetic tree of Athila elements in *G. urbanum* and *G. rivale* is shown to the left. To the right of the tree, two bar plots are shaded to indicate which elements belong to *G. urbanum* (right) and *G. rivale* (left), with coloured red and blue shading indicating elements found within the centromeres. To the right, for each chromosome of each species, coloured dots indicate which tree tip element is located on which chromosome, with darker shading indicating centromeric copies. No bias for centromere-specific clustering can be observed. The phylogenetic tree was constructed using concatenated core domains of five transposon genes **e.** Sequence identity histograms generated by pairwise comparison of LTR sequences (left) and internal domain (right) of all Athila elements found within centromere boundaries for *G. urbanum* and *G. rivale*. High identity in the between-species comparison suggests the presence of a single Athila family in the centromeres of both species.

**Extended Data Figure 13.**
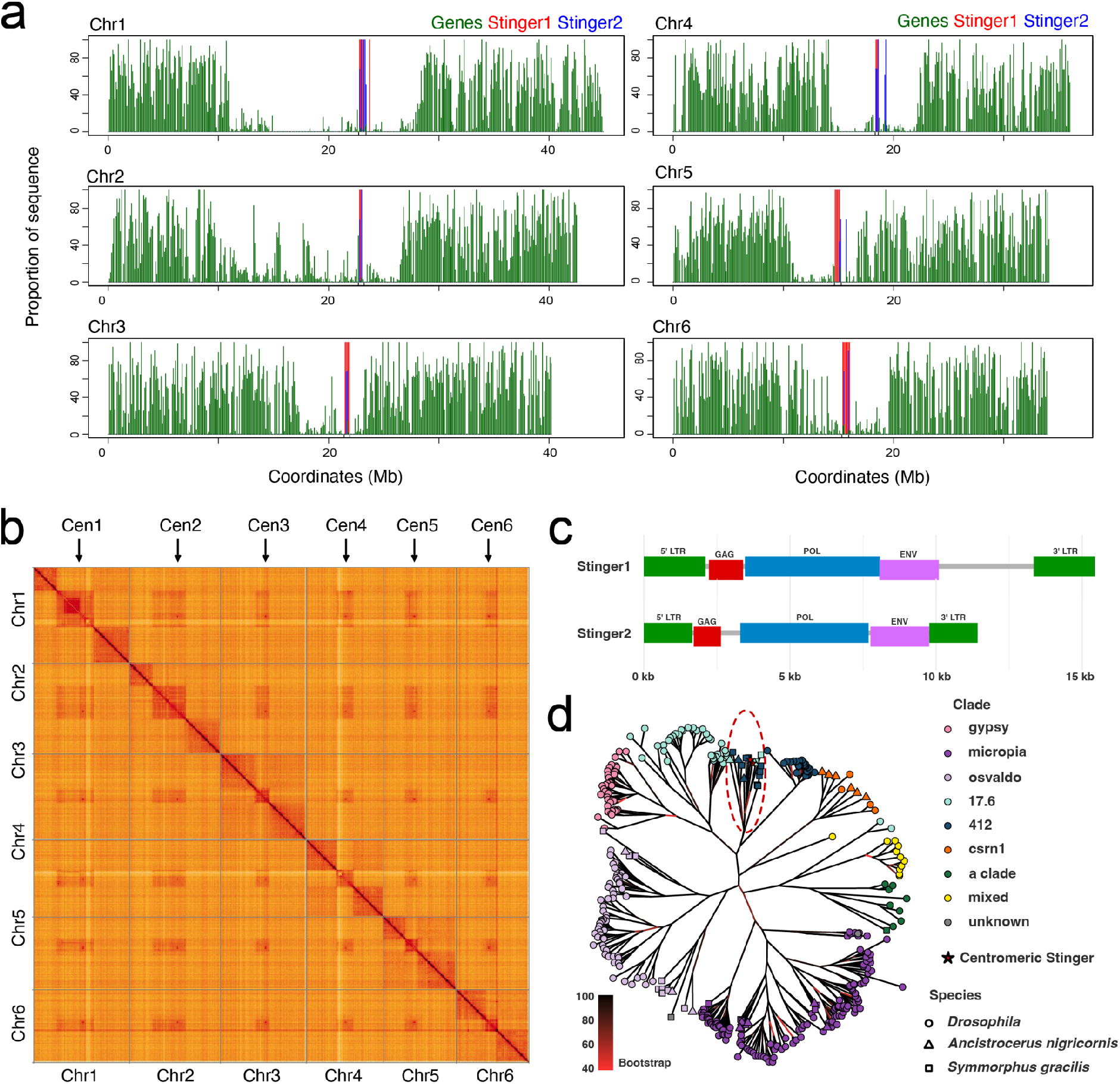
*Ancistrocerus* Gypsy LTR centrophilic transposons. **a.** Plots of the six *Ancistrocerus nigricornis* chromosomes showing the proportion of annotation that corresponds to genes (green), Stinger1 (red) and Stinger2 (blue) family Gypsy LTR retrotransposons. **b.** A Hi-C contact matrix of the *Ancistrocerus nigricornis* chromosomes with the location of the Stinger clusters indicated by black arrows. **c.** Diagrams showing the structure and composition of the two centrophilic Stinger transposon families we identified. **d.** A phylogenetic tree inferred from an alignment of the largest ORF from 43 Gypsy families from *Ancistrocerus nigricornis*, including the two centromeric families, 18 families from *Symmorphus gracilis* (wasp) and 329 Gypsy families from 30 different species of *Drosophila*. The tip colors correspond to the TEsorter classification by Gypsy clades. The tip shape follows the species of the family (a single shape is assigned for all *Drosophila* transposons). Branches leading to nodes are colored by their bootstrap values. Gypsy elements, including Stinger, from both wasp species form a monophyletic clade, similar to clades 17.6 and 412 from *Drosophila* (red oval).

**Extended Data Figure 14.**
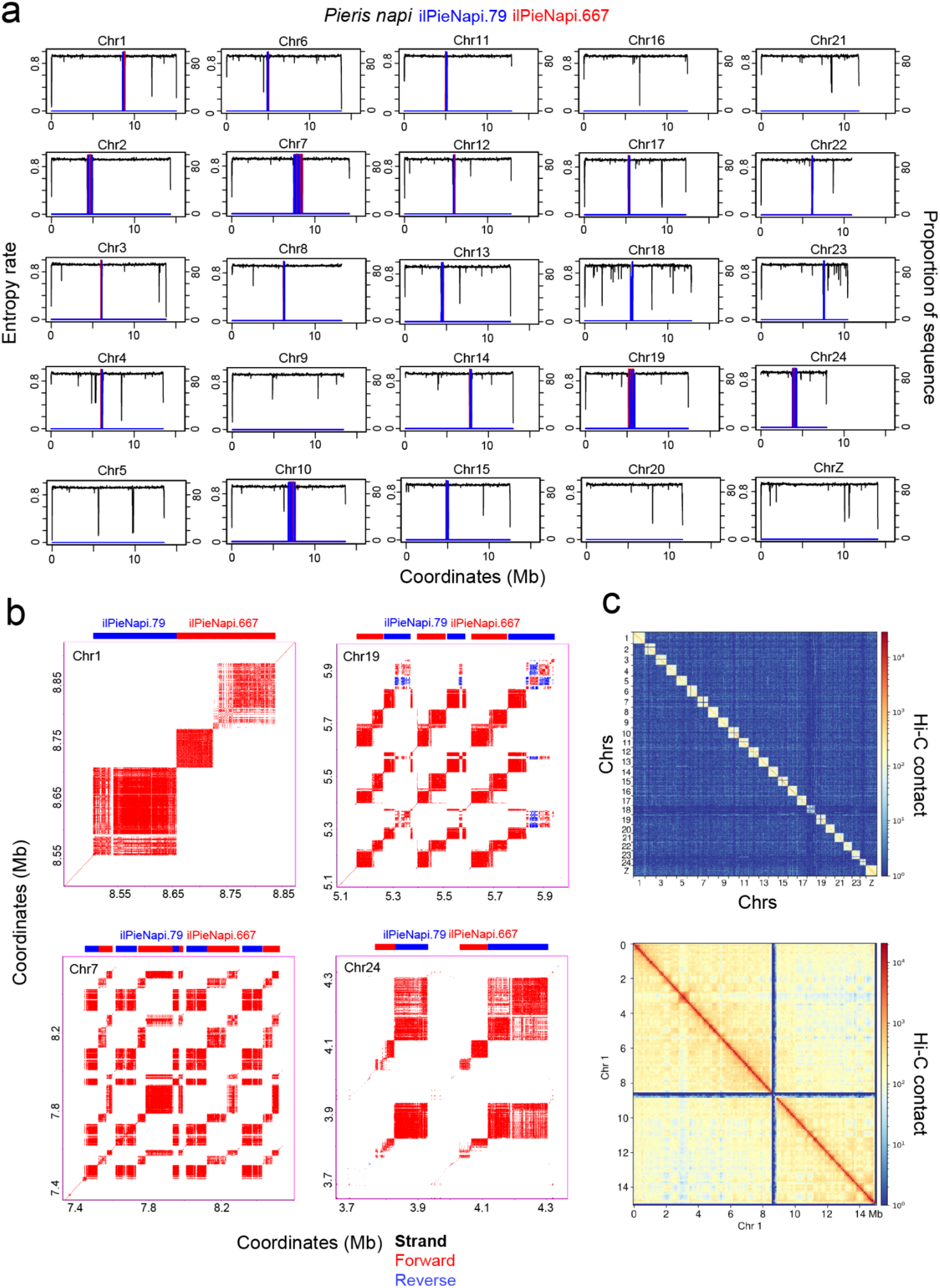
Satellite arrays in the holocentric butterfly *Pieris napi.* **A.** Plots of the *P. napi* chromosomes showing entropy rate (black) and the proportion of the sequence corresponding to the satellite repeat families ilPieNapi.79 (blue) and ilPieNapi.667 (red). **b.** Sequence identity dot plots of representative satellite array regions from *P. napi* chromosomes, where red and blue shading indicate forward and reverse strand sequence similarity. At the top of each dot plot, coloured blocks indicate the location of ilPieNapi.79 (blue) and ilPieNapi.667 (red) repeat arrays. **c.** Heat maps showing Hi-C contact probability (Hi-C resolution was 100 kb) within and between chromosomes in holocentric *P. napi* (Lepidoptera, Arthropoda), showing a pattern typical of holocentric genomes, despite the presence of satellite repeat arrays that resemble those observed in monocentric genomes.

**Supplementary Table 1. A genomic atlas of eukaryotic centromere architecture.** The table provides species names, three levels of taxonomic classification, the CAP centromere architecture state, the Darwin Tree of Life ID (TolID), the GenBank assembly accession, the number of chromosomes, their total length in base pairs and %GC content, the number of scaffolds in the assembly, the length of the scaffolds in base pairs, the Scaffold N50, and the contig N50. Of these 325 species, the following have previously had centromeres characterised; *Arabidopsis thaliana*^30^, *Luzula sylvatica*^40^, *Gallus gallus*^115^, *Rosa canina*^41^, and *Taeniopygia guttata*^41,116^.

**Supplementary Table 2. Centromere-associated satellite repeats from across eukaryotes.** For each satellite family identified as centromere candidates, we list the species of origin and taxonomic information, the satellite name (derived from the species TolID and the mean length of the satellite family), the count of repeats in the genome, their mean length in base pairs, their mean GC content, the number of host genome chromosomes, the length of the chromosomes and their GC% content.

**Supplementary Table 3. Transposon content of centromeric satellite arrays.** For species with CAP-predicted satellite arrays, the table contains analysis as directed in column ‘analysis_step’; a value of 1 indicates the total centromere length; a value of 2 shows the total length of the satellite arrays (listed as ‘TR_nt_’ in column ‘annotation_feature’), the length of non-satellite interruptions in the arrays (‘interruptions_nt_’ in column ‘annotation_feature’), and their proportion of the total centromere length (column ‘centromere_prop’); a value of 3 resolves the non-satellite interruptions by showing the total length of transposons (‘TE_nt_’ in column ‘annotation_feature’) and any other low-copy satellites that have not been classified as part of the centromeric arrays (‘otherTR_nt_’ in column ‘annotation_feature’), as well as their proportion of the total centromere length; a value of 4 resolves transposons (‘TE_nt_’) into Class_I_LTR (LTR retrotransposons and retroviruses), Class_I_TPRT (non-LTR retrotransposons such as LINEs that undergo target-primed reverse transcription), Class_I_other (Penelope and Tyrosine Recombinase elements), and Class_II_DNA (elements that have terminal inverted repeats such as Mutator and CACTA, and also Helitrons). ‘TE_unclass’ and ‘repeat_region’ refer to annotations of the EDTA pipeline that were not reassigned to a specific transposon class by our dedicated reannotation pipeline (see Methods); finally, a value of 5 further resolves transposons into their superfamily phylogenetic level.

**Supplementary Table 4. FISH probe sequences.** The DNA sequence of FISH probes used for cytogenetic detection of centromere sequences are provided from 5′ to 3′. For species where we used oligo-pools for FISH probes, the sequences are provided on the project Github page (https://github.com/vlothec/eukaryotic_centromere_architecture).

## Materials and Methods

### Darwin Tree of Life chromosome-level genome assemblies

All DToL genome assemblies were generated using a combination of Pacific Biosciences (Pacbio HiFi) long reads and Hi-C Illumina reads^117^. Pacbio HiFi reads are the input for k-mer analyses and *de novo* genome assembly, and Hi-C reads are used for scaffolding and manual curation^117^. Assembly decontamination and manual curation followed the methods described^117^. All assemblies are publicly available under the European Nucleotide Archive project accession PRJEB40665. **Supplementary Table 1** presents NCBI accession numbers (e.g. GCA_933210815.1) for each assembly, as well as chromosome number and length, scaffold number and length, and scaffold and contig N50. Where available, a link to the associated DToL genome note is provided in **Supplementary Table 1**.

### Annotation of tandem repeats

For satellite repeat annotation we applied TRASH, which uses periodic k-mers to identify tandem repeats^46^, using an updated version developed for this study (github.com/vlothec/TRASH_2). For each genome, two runs were performed; one using an upper repeat detection limit of 1 kb, which identified most centromeric satellites, and a second run with an upper repeat detection limit of 10 kb and a lower limit of 1 kb. We applied the longer repeat detection window to genomes where entropy rate analysis indicated the presence of tandemly repeated sequences that were not identified by TRASH using the upper detection limit of 1 kb. The identified tandem repeat families were refined by removing dispersed repeats that had a similarity score against the family consensus lower than 40%, or which appeared in contiguous repeat arrays shorter than 1 kb. Centromeric arrays were defined as clusters of arrays of CAP-identified centromeric satellite families that were interrupted by no more than 100 kb of non-satellite sequence. A 50 kb limit was used for holocentric species with periodic satellite arrays.

### Annotation of transposons

Transposons were identified using the EDTA^45^ pipeline (v2.0.1) with parameters --anno 1 and --sensitive 1, and lineage-specific libraries from Repbase^118^ as --curatedlib (e.g. dcotrep.ref for eudicot plants). The Repbase libraries used are available here: https://www.girinst.org/server/archive/RepBase28.06/. We filtered the initial EDTA annotation by intersecting transposon annotation with satellite repeat coordinates from TRASH. Transposons that overlapped >80% of their length with satellites were removed. We then parsed EDTA annotations that were classified as ‘repeat_region’ with the repeat_region_rescue script to reassign them to specific transposon types (repeat_region_rescue script is available at https://github.com/vlothec/eukaryotic_centromere_architecture). This rescue pipeline combines tables of transposon family names retrieved from Dfam (https://www.dfam.org/classification/tree), the EDTA sequence ontology, and the headers of the input libraries, which were then used in sequential steps to match the ‘Name’ and ‘Classification’ information in the attribute column of the output gff3 file. As an additional step for plant species, LTR retrotransposons in the reassigned gff3 file were further classified by TEsorter^48^ (v1.4.6) into lineages using parameters -db rexdb-plant -nolib. The filtered and reassigned files were used as input for the centromere annotation pipeline (CAP), for annotation of interruptions to the satellite arrays, and other downstream analysis. Scripts associated with this transposon analysis are located in the te_and_interruption_analysis folder at https://github.com/vlothec/eukaryotic_centromere_architecture.

Phylogenetic analysis of the Athila LTR retrotransposon lineage in plants was conducted by parsing the EDTA gff3 files for structurally intact LTR retrotransposons and using TEsorter to identify candidate intact Athila elements. We retained Athila elements that contained all five main genes (gag, protease, reverse transcriptase, RNAseH, and integrase) based on the hidden Markov model analysis of TEsorter, concatenated their amino acid sequences, aligned them using MAFFT^119^ (v.7.490, --retree 2 --maxiterate 100), and generated a maximum-likelihood tree using FastTree^120^ with default parameters. SoloLTRs were identified using soloLTRseeker (https://github.com/estpr/soloLTRseeker). In brief, soloLTRseeker generates a non-redundant high-quality library using the LTRs of structurally intact retrotransposons that were classified in advance into LTR lineages by TEsorter. Using this library as a query, BLASTn scans the genome for putative soloLTRs with thresholds of 99% query coverage, and 80% sequence identity. The strict query coverage threshold ensures that we have identified nearly the entire length of the input LTRs in the genome. Flanking 200 bp sequences of each putative soloLTR locus were compared with sequences of the same length from both termini of the internal domain of intact elements from the same lineage as the best BLASTn hit. We use an 80% similarity threshold to identify and filter out cases of partially deleted transposons that are not soloLTRs. Upstream and downstream sequences of 6 bp length from each soloLTR were locally aligned to annotate target site duplications (TSDs).

We analysed in detail the content of the part of satellite centromere arrays that was not covered by the TRASH-defined tandem repeats. We termed these regions ‘interruptions’. Our approach was to use the transposon annotation to calculate the transposon content of interruptions, and to classify them according to their transposon profile. For example, one interruption may contain two types of transposon annotation (e.g. LTR retrotransposons and Class II DNA transposons), whereas another interruption may contain only non-LTR retrotransposon annotation. This classification system grouped interruptions into distinct transposon categories, which allowed us to visualise and compare their content, sequence similarity, and position along centromeres, within and between categories.

### Analysis of transposon-state centromeres

In the transposon-based centromeres of *G. urbanum* and satellite-based centromeres of *G. rivale*, we used as queries intact Athila elements to run BLASTn against the parts of the centromeres that were not identified as transposons or satellites by our annotation pipeline (mostly short interruptions). When the coverage of the interruptions was greater than 80%, the entire interruption was reassigned as Athila. For *G. urbanum*, we reconstructed older strata of intact Athila transposons by computationally removing intact elements, patching their flanking sequences, and running EDTA as in the original genomes.

As an additional approach to resolve the internal nesting structure of transposable elements within candidate centromere regions, we developed an iterative excision procedure that reconstructs layer by layer the order in which nested transposons are inserted into a given locus. For each region of interest, intact transposons annotated by EDTA were first excised as layer 0, and the flanking sequence was stitched together to yield a de-nested current sequence, which was then used as the BLASTn (v2.12.0) target for a self-referential search, using the full set of intact transposons as query sequences. Searches begin at a nucleotide identity threshold of 90% and decrease in steps of 5% to a floor of 80%, with only alignments covering at least 90% of the query length retained as candidate insertions. This cycle continues until the identity floor is reached, or no further transposons can be excised. The number of iterative excision layers required to fully resolve a region was regarded as its relative nesting depth.

Transposable elements in *Ancistrocerus nigricornis* were identified using EDTA^45^ and a development version of pantera^121^, based on FastGA^122^. The identified *A. nigricornis* Gypsy families were classified in different clades using TEsorter^48^ with parameters -db gydb. We extracted the largest ORF from each family and confirmed they encoded the POL polyprotein. We aligned the ORF sequences using MAFFT^119^ with options --localpair --maxiterate 1000 -- thread 8, together with POL ORFs found in Gypsy elements from the wasp *Symmorphus gracilis* (a member of the Eumeninae that is ∼20 million years diverged from *A. nigricornis*) and from 30 species of *Drosophila* obtained using the same development version of pantera, together with curated Gypsy sequences available in Flybase. The resulting alignment was used to produce an unrooted phylogenetic tree using IQ-TREE 3^65^ with options iqtree3 -m Q.INSECT+R8 -B 1000 -T AUTO.

### Annotation of protein-coding genes

To identify protein-coding genes *de novo* we used Helixer^47^. Helixer is a machine learning-based tool for *ab initio* gene prediction in the absence of experimental data such as RNA sequencing^47^. Helixer was run in the matched lineage-specific mode for the genome being analysed.

### Annotation-free estimates of sequence repetition

The context-tree weighting algorithm (CTW) is a general modelling method, originally introduced for data compression^49,50^. For discrete symbol sequences with relatively small alphabets, such as DNA sequences, it provides an effective tool for estimating the entropy rate of the underlying data generating mechanism^49,50^. The CTW algorithm may be viewed as a computationally efficient method for calculating the marginal likelihood of observed sequences, based on a hierarchical Bayesian model that captures dependence in an effective and parsimonious manner. This marginal likelihood has been shown to approximate the true log-likelihood of the data, with respect to its true distribution. This guarantees that the resulting entropy estimates are consistent, and that both their bias and variance are minimal. For each genome, we coded chromosome sequences as 0, 1, 2, and 3, replacing A, T, G, and C, in adjacent 10, 50, or 100 kb windows, depending on genome size, and applied the CTW function from the R package BCT^50^, with a depth parameter of 10. CTW values obtained were multiplied by the base-2 logarithm of 1 over e (-log2e), and divided by window length, minus the depth value. The resulting entropy rate measurements were then plotted along chromosomes, and compared to gene, tandem repeat, and transposon annotation to help identify candidate centromere regions.

### Centromere Analysis Pipeline (CAP)

CAP takes as an input the genomic assembly and annotation files of tandem repeats, transposons, and genes (**Extended Data Figure 1**). We used TRASH, EDTA, and Helixer outputs as these input files, respectively, generated as described in the annotation methods. CAP identifies the most abundant tandem repeats per chromosome, and within the genome, and calculates the mean pairwise similarity (Levenshtein distance normalised by mean repeat length) as a measure of repeat homogeneity, as well as consistency of repeat monomer size (repeat length distribution standard deviation normalised by mean repeat length). For each chromosome, in each genome, three plot panels are generated. The upper panel contains details of the identified satellite families. The middle plot shows %GC content in 2 kb windows, and CTW entropy rate in 100 kb windows. In addition, tandem repeat density is plotted in 10 kb windows (grey histogram bars), with satellite families from the table above plotted independently with colour-coding matching the table text. The lower plot shows transposon density in 100 kb windows (grey histogram lines), with individual transposon classes traced in colours that match the global transposon legend. The two trendlines on the lower plot are moving averages of gene content (in 50 bins, green) and combined transposon and tandem repeat content (in 50 bins, blue). The trendlines are useful to visually assess regions of increased repeat and decreased gene content, which is characteristic of centromeric and pericentromeric regions. CAP is available on GitHub (https://github.com/vlothec/CAP).

We profiled centromere architecture of each analysed genome, identifying satellite families and/or transposon lineages occupying putative centromeric regions using CAP. Each centromeric satellite family classified by CAP was named using the species Tree of Life Identifier (ToLID) combined with the repeat mean length (**Supplementary Table 2**). For species that did not exhibit high-scoring satellite candidates, we used the same outputs to search for regions of high transposon density and low gene exon density. When the transposons were of a consistent class between chromosomes, we classified the genome as monocentric transposon architecture. A subset of species was classified as showing a mixed architecture, where some chromosomes possessed high-scoring satellite arrays, and others possessed transposon clusters. In addition, Hi-C contact maps were used to identify centromeres in fungal genomes as regions with reduced intra-chromosomal contact with the chromosome arms and increased inter-chromosomal centromere-centromere interactions^55,56^.

For centromere boundary definition in satellite-architecture genomes, in the first step, arrays were mapped as stretches of tandem repeats with interruptions of less than 100 kb between them. The centromere regions were defined by the arrays of the candidate centromeric satellite family. In some cases, dispersed centromeric repeats could be found at low copy elsewhere on the chromosome. Therefore, we applied a filter which excluded arrays where, (i) an array was fully contained within the first or last 25% of all repeats of that family along the chromosome, and (ii) an array accounted for less than 10% of all repeats of that family on the chromosome. In holocentric species with satellite arrays, the maximum interruption used was 50 kb, and all arrays were considered centromeric. Centromere boundary definition for transposon-based architectures was determined through visual inspection of sequence identity dot plots to identify regions of elevated transposon enrichment.

For CAP benchmarking and validation, the following genomes and CENP-A/CENH3 ChIP-seq and input datasets were analysed; (i) *Homo sapiens* CENP-A ChIP-seq (SRX255043) and input (SRX255044) data were aligned against the CHM13 (GCF_009914755.1) assembly^31^; (ii) *Mus musculus* CENP-A ChIP-seq (SRX20577194) and input (SRX20577193) data were aligned against the GCA_000001635.9 assembly^58^; (iii) *Timema douglasi* CENP-A ChIP-seq (SRX28824591) and input (SRX28824592) data were aligned against the GCA_040436065.2 assembly^61^; (iv) *Oryza sativa* CENH3 ChIP-seq (SRX27798377) and input (SRX27798382) data were aligned against the GCA_001623365.2 assembly^59^; (v) *Zea mays* CENH3 ChIP-seq (SRX17512265) and input (SRX17512267) data were aligned against the GCA_022117705.1 assembly^60^; (vi) *Triticum monococcum* CENH3 ChIP-seq (DRX799804) and input (DRX799805) data were aligned against the GCA_057380995.1 assembly^12^; (vii) *Pisum sativum* CENH3 ChIP-seq (ERR9981080) and input (ERR9981081) data were aligned against the GCA_977071245.1 assembly^114^; and (viii) *Cryptococcus deuterogattii* R265 CENP-A ChIP-seq (SRX3049403) and input (SRX3049404) data were aligned against the GCA_002954075.1 assembly^33^. In each case we followed our CAP workflow and assessed how our centromere classifications related to the known CENP-A/CENH3-occupied sequences.

To map CENP-A/CENH3 across the benchmark genome assemblies, input and CENP-A ChIP-seq reads were aligned to each reference assembly using Bowtie2 with default settings. Reads were mapped as paired-end, or single-end, depending on the structure of the underlying sequencing library. The resulting read alignments were coordinate sorted with SAMtools. For each assembly, genomic bins of fixed width were generated. Per-bin read counts were computed separately for the ChIP and input alignments using BEDTools coverage (-counts). To allow direct comparison of enrichment magnitude across libraries of different sequencing depth, per-bin counts were converted to reads-per-million (RPM) using the total number of mapped reads for each BAM file (obtained via samtools idxstats, summed across all reference sequences) as the normalization denominator. Bins with zero coverage were assigned a pseudocount of 0.5 prior to ratio calculation to avoid undefined values. Relative CENP-A/CENH3 enrichment per bin was then calculated as log2(ChIP/input). Per-bin log2 enrichment ratio was plotted along chromosomes using R ggplot2.

### Species phylogenetic tree building and dating

A maximum-likelihood species tree was inferred from the 325 DToL Helixer-predicted proteomes using FastSpeciesTree^64^ (v1.0). FastSpeciesTree performs ortholog identification, alignment, and tree inference end-to-end and used DIAMOND^123^ blastp to perform homology search of each proteome against the eukaryota_odb10 BUSCO reference orthologs. For each single-copy ortholog, the best-hit local alignment is anchored to the reference-query coordinates and gap-padded to the query length. Then, these ortholog alignments are concatenated into a query-anchored pseudo-alignment. Orthologs that were absent from a proteome were encoded as gaps. Genes present in at least 80% of species (no more than 20% missing taxa) were retained, yielding 194 single-copy loci. FastSpeciesTree internal trimming retained only parsimony-informative columns; that is, positions with at least two amino-acid states in >1 taxa (gaps and ambiguous residues were ignored), discarding constant and singleton sites and any locus with fewer than five informative columns. The supermatrix comprised 52,754 amino acid sites, with 4.2% missing data. A maximum-likelihood tree was inferred using the supermatrix with IQ-TREE^124^ (v2.3.4) using ModelFinder^125^ to select the best-fit substitution model independently per gene partition. This was selected from models LG, JTT, Q.INSECT, Q.YEAST, Q.BIRD, Q.MAMMAL, and Q.PLANT, together with +I, +G and +I+G rate-heterogeneity models. Q.INSECT and LG models were selected for the large majority of partitions. Branch support was assessed using 1,000 ultrafast bootstrap replicates (UFBoot) and 1,000 SH-aLRT replicates.

The tree was midpoint-rooted, and calibration nodes were placed at well-supported clades, and each was assigned a minimum-maximum age constraint. Sixty-two constraints were used in total, consisting of one root constraint (Eukaryota, 1,085-1,671 Mya), and 61 internal nodes distributed across Opisthokonta root (n=1), Metazoa (n=31), Viridiplantae (n=21), and Fungi (n=8). The bounds were taken from the corresponding TimeTree 5^67^ divergence-time confidence intervals, retrieved automatically using PAReTT (https://github.com/LSLeClercq/PAReTT) for 58 nodes. The remaining 4 nodes, which lacked a reported TimeTree range, were bounded by the TimeTree median ±20%. The inferred tree topology was time-calibrated using the *chronos* function from the ape R package^66^.

To select a dating model, we compared a range of penalised-likelihood models and benchmarked each against TimeTree by MRCA node-age concordance. Trees were pruned to the 210 species shared with TimeTree, and the age of every species pair’s MRCA was computed on each tree. The resulting 21,945 pairwise comparisons (all pairs among the 210 shared species) were grouped by their TimeTree divergence into 213 distinct node-age comparisons, and concordance was summarised by Pearson’s *r* and *R²* between our estimate, and the TimeTree-derived ages. We tested correlated, relaxed, and discrete rate models, each across the smoothing parameter values λ=0, 0.1, 1, and 10. The correlated-rates model with λ=0.1 gave the closest agreement with TimeTree node ages (Pearson’s *r*=0.98, *R²*=0.97) and used for further analyses.

### Ancestral state reconstruction modelling of centromere state evolution

Ancestral state reconstruction (ASR) was performed using Markov k-state (Mk) continuous-time models fitted by maximum likelihood using *fitMk* from the phytools R package^68^. This approach estimates transition rates between discrete character states and reconstructs marginal state probabilities at all internal nodes of the phylogeny^68^. Using the CAP centromere architecture states on the species tree, we used phytools to compare an equal-rates (ER) model, a symmetric (SYM) model, an all-rates-different (ARD) model, and several custom directional models. All models were evaluated using the Akaike Information Criterion (AIC and cAIC), and Akaike weights. ASR was performed independently on three datasets: the full 325-species tree, a Metazoa subtree, and a Viridiplantae subtree. Seven models were evaluated per tree, varying in the number and structure of permitted transition rates, as well as to test irreversibility of the holocentric state (**Extended Data Fig. 5**). The mixed satellite/transposon centromere and cryptic states was observed in too few to support reliable rate estimations, and so both categories were pruned from the tree prior to fitting, leaving a 3-state alphabet of holocentric, satellite, and transposon states, and 277 tips for the full tree. Ancestral states at internal nodes were reconstructed under the best-supported model (All-Rates-Different with Holocentric irreversible; ARD_irrevH).

### CENH3/CENP-A sequence curation and analysis

CENP-A/CENH3 protein sequences were retrieved from the DToL genomes using a two-stage homology-search strategy combining DIAMOND searches, with genome-based rescue for species where DIAMOND was insufficient; followed by phylogeny-guided manual curation. In the first stage, Helixer-predicted proteomes for all 325 species were searched against a curated database of 450 CENP-A/CENH3 reference sequences compiled from a published eukaryotic CENH3 database^3^, and KEGG Orthology entries K11495 (CENP-A) and K11253 (CENH3) using DIAMOND^123^ (v.2.1.11) in sensitive mode. Hits were retained if they matched a CENP-A-specific HMM profile using HMMER^126^ (v3.4), which yielded 7,846 candidate sequences from 253 species. Reciprocal best-hit analysis was used to independently confirm CENP-A ortholog presence in these 253 species. In a second stage, for species where no CENP-A candidate was detected using DIAMOND, a genome-level rescue was additionally performed using miniprot^127^ (v0.15), aligning the references directly to the genome assemblies. After collapsing overlapping predictions within 1 kb windows, 1,299 of 1,315 candidates (98.8%) passed the CENP-A HMM filter, extending CENP-A/CENH3 detection to a further 21 species. The full CENP-A/CENH3 candidate ortholog pool was combined with five well-characterised CENP-A/CENH3 reference sequences from *Homo sapiens* (UniProt P49450), *Mus musculus* (O35216), *Danio rerio* (Q803H4), *Drosophila melanogaster* (Q9V6Q2, *cid*), and *Caenorhabditis elegans* (P34470, *hcp-3*). This combined pool was aligned with MAFFT^119^ (v7.526), trimmed with ClipKit^128^ (v2.3.0), and a preliminary maximum-likelihood tree was inferred with FastTree^120^ (v2.1.11) to guide manual curation. Sequences forming a monophyletic clade with the five CENP-A reference orthologs, clearly separated from the H3 outgroup, were retained as *bona fide* CENP-A/CENH3 orthologs.

Canonical histone H3-like sequences were retrieved from the same Helixer proteomes by HMMER^126^ (v3.4) hmmsearch using a histone H3-specific HMM profile. We retained up to three top-scoring sequences per species, yielding 901 sequences in total. Ten archaeal histone sequences were included as a phylogenetic outgroup^129^. The curated CENP-A, H3-like and histone archaeal sequences^129^ (OLS22332.1, OLS24873.1, OLS21974.1, KKK41979.1, KXH71038.1, OLS18261.1, OLS16336.1, BAD86478.1, OIO61677.1, and OIO41945.1) were aligned with MAFFT^119^, trimmed with ClipKit^128^, and a final maximum-likelihood tree was inferred with IQ-TREE2^65^ (v2.3.4) using the command iqtree2 -m MFP -B 1000 -bnni -nm 2000. The substitution model VT+G4 was selected by ModelFinder^65^ based on the BIC criterion. The tree search converged at iteration 404. Applying the same criterion as for the preliminary tree (i.e. retaining sequences forming a clade with the characterised CENP-A sequences) manual curation was performed yielding a final set of 422 CENP-A/CENH3 sequences, and 897 H3-like sequences that were used for subsequent analyses.

Shannon’s entropy was computed per alignment position independently for CENP-A/CENH3 (n=422) and H3-like (n=897) sequences from the combined sequence alignment used for phylogenetic inference, following^130^. For each alignment position i, Shannon’s entropy was calculated as H = –Σ(pi × log2(pi)) and normalized by dividing the calculated entropy by the maximum Shannon’s entropy (4.32 for amino acids, meaning all 20 standard amino acids are equally present), where pi represents the frequency of each amino acid at that position. Columns where more than 85% of sequences within a given group had a gap were masked, per group. Entropy values were normalised between 0 and 1. The resulting per-group entropy profiles were overlaid with the predicted protein secondary structure of *Arabidopsis thaliana* CENH3 (UniProt Q8RVQ9). Histone α-helix positions obtained via STRIDE^78^ were projected onto the trimmed alignment coordinates, allowing positions in CENP-A to be contextualised within the histone fold domain, the CENP-A targeting domain (CATD), and the N- and C-terminal domains.

Positions in the CENP-A/CENH3 alignment that differ systematically between functional groups (specificity-determining positions, SDPs) were identified using GroupSim^75^, which scores each alignment column by the difference between within-group and between-group residue similarity, favouring columns conserved within each group but divergent between them. GroupSim was run on a trimmed alignment from which columns with >85% gaps across all sequences had been removed. Gaps were treated as a 21st character, and each column score was smoothed over a ±3-residue window in which neighbouring columns were weighted (λ=0.7) by their per-column conservation (Jensen–Shannon divergence), and scores were normalised between 0 and 1. Two comparisons were performed using GroupSim: (i) CENP-A vs histone H3, contrasting 422 CENP-A/CENH3 sequences with 897 H3-like from the alignment, and (ii) Satellite vs Transposon, comparing CENP-A/CENH3 sequences from satellite-state centromere genomes, compared against those with transposon-state centromeres. To ensure SDPs were not driven by uneven phylogenetic sampling, the specificity score was recomputed with a clade-level weighting scheme in which each sequence was weighted by 1/n_clade, with n_clade being the number of sequences from the same broad taxonomic group (insects, vertebrates, Viridiplantae, fungi, and other invertebrates) within its comparison group, and weights normalised to a mean of 1 per group. Positions with a clade-weighted z-score ≥2 were taken as significant SDPs.

### Fluorescent *in situ* hybridization and immunocytology

FISH probe design was performed using the available genome annotation, in addition to using independent methods for transposon and tandem repeat annotation; DANTE/DANTE-LTR^131^ and TideCluster (https://github.com/kavonrtep/TideCluster), respectively, using default settings. For immunocytogenetic analysis *Ailanthus altissima* seeds were collected from a tree in Prague, Czech Republic (GPS coordinates: 50.0995028N, 14.3805733E); *Ballota nigra* plants were collected in Hluboká nad Vltavou, Czech Republic (GPS coordinates: 49.0492906N, 14.4427019E); *Chamaenerion angustifolium* plants were collected in České Budějovice, Czech Republic. (GPS coordinates: 49.00964N, 14.44140E); *Geum urbanum* seeds were collected from a plant in České Budějovice, Czech Republic (GPS coordinates: 49.00979N, 14.44253E); *Linaria vulgaris* plants were collected in České Budějovice, Czech Republic (GPS coordinates: 49.00737N, 14.44072E); *Mercurialis annua* plants were collected and provided by Jindrich Chrtek (Institute of Botany, Pruhonice, Czech Republic); and *Solanum dulcamara* plants were collected in Malé Chrášťany, Czech Republic (GPS coordinates: 49.0390608N, 14.3089522E).

To analyse metaphase stage cells, fresh root tips were immersed in ice-cold water at 4°C for 24 hours. The next day, the material was fixed in Carnoy’s solution (3:1 ethanol: acetic acid) overnight at room temperature and then stored at -20°C. Root tips were washed twice in cold distilled water, and twice in Tris buffer, each for 5 minutes. Meristems were digested with 2% cellulase, 1% pectinase and 0.2% pectolyase in citrate buffer at 37°C, with digestion times adjusted for each species (*A. altissima* 120 minutes, *B. nigra* 60 minutes, *C. angustifolium* 60 minutes, *G. urbanum* 75 minutes, *L. vulgaris* 50 minutes, *M. annua* 60 minutes and *S. dulcamara* 50 minutes). Chromosome spreads were prepared using the air-dropping method. FISH was performed using oligonucleotide probes synthesized by Integrated DNA Technologies (*G. urbanum* and *L. vulgaris*), or by Eurofins Genomics (*B. nigra*, *M. annua*, and *C. angustifolium*), following^39^. Chromosomes were denatured in 30 μL of hybridization mixture (50% formamide, 10% dextran sulfate, 2×SSC, and 0.5 μM probe labeled with biotin during synthesis). The denaturation time was adjusted for each species (*B. nigra* 1 minute 20 seconds, *C. angustifolium* 1 minute 10 seconds, *G. urbanum* 2 minutes 30 seconds, *L. vulgaris* 1 minute 25 seconds, *M. annua* 40 seconds, and *S. dulcamara* 1 minute 40 seconds). The hybridization and washing temperatures were adjusted to account for varying AT:GC content between species, with 37°C for hybridization and 42°C for washing being the default. Post-hybridization washes were conducted in 0.1×SSC and probes were detected using Streptavidin-Alexa Fluor 568 (Thermo Fisher Scientific). For *S. dulcamara* and *A. altissima*, due to sequence complexity and relatively low transposon density in the centromeres, we used oligo-pool probes synthesized by Integrated DNA Technologies. The FISH procedure was performed using 25 pmol of the oligo-pool probe per slide, 32°C for hybridization, and 37°C for washings. Chromosomes from *M. annua* previously hybridized with the ddMerAnnua.93 probe were re-hybridized with the ddMerAnnua.62 probe. Slides were placed in a humid chamber at 37°C for 10 minutes to reduce Vectashield viscosity. After removing the coverslips, slides were washed as follows: twice for 30 minutes each in 4×SSC+0.2% Tween at room temperature; twice for 5 minutes each in 2×SSC at RT; once for 10 minutes in 50% formamide/2×SSC at 55°C; and finally, twice for 5 minutes each in 2×SSC at room temperature. Slides were dehydrated in 70% and 90% ethanol for 5 minutes each, air-dried, and used for a new FISH procedure with the ddMerAnnua.62 probe. All FISH probe sequences are provided in **Supplementary Table 4**.

For immunostaining, fresh root tips were fixed in 3% formaldehyde diluted in Tris buffer (10 mM Tris, 10 mM Na₂EDTA, 100 mM NaCl, pH 7.5) at 4°C for 30 minutes, with the first 5 minutes under vacuum. Samples were then washed in Tris buffer on ice for 30 minutes. Chromosome suspensions were prepared in LB01 buffer^57^. To visualize functional centromeres, we used an affinity-purified rabbit polyclonal antibody against the kinetochore protein KNL1^57^. Immunostaining was performed, as described^57^. An exception in chromosome preparation was made for *B. nigra*, where meristem fixation and chromosome preparation were performed as described for FISH to enhance chromosome spreading, as KNL1 immunostaining performed well in material fixed in Carnoy’s solution (3:1 ethanol:acetic acid) for this species. For combined detection of KNL1 and centromeric probes in *A. altissima* and *B. nigra*, chromosomes were prepared as for *in situ* immunostaining. Immunodetection was followed by FISH and both procedures were carried out as described above. After immunostaining, and/or FISH preparation, the slides were post-fixed in 4% formaldehyde diluted in 1×PBS for 10 minutes at room temperature, chromosomes were counterstained with 4,6-diamino-2-phenylindole (DAPI) and slides were mounted with Vectashield mounting medium (Vector Laboratories). A Zeiss AxioImager.Z2 microscope equipped with an AxioCam 506 mono-color camera, and with an Apotome 2.0 device was used. Image acquisition was performed using the software ZEN 3.2 blue edition (Carl Zeiss GmbH).

### Satellite repeat analysis

To investigate sequence similarity across satellite families we took a k-mer based approach. To decide on the most reasonable *k* length for this dataset, we performed a sweep of length values to investigate the number of possible versus observed k-mers. We selected a *k* of 17 as the number of observed k-mers was substantially less than the total possible for this value. We generated 17-mers from all copies of each satellite family and conducted pairwise comparisons of k-mers between families, including randomized sequences with varying GC content as controls. Satellite families with mean lengths shorter than 17 base pairs long were excluded from this analysis. For comparison, random sequences were generated with the python random package with varying GC content and varying sequence length. K-mer comparison was performed with mash, using the following command and using the outputted shared k-mer counts:

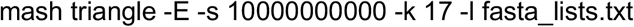

To further analyse satellite repeat sequence similarity and how it decays with species divergence time, for each family, a random 10% sub-sample of its monomers (restricted to ≥50 bp) were drawn, capping at 1,000 sequences per family, yielding 148,794 sequences in total. For every pairwise combination of satellites, sequences were compared using local alignment with BLASTN^132^ (v2.16.0+), following the approach in Melters et al^6^. As tandem-repeat monomer boundaries are arbitrary, each query set was searched against a database in which every monomer was represented as a heat-to-tail tandem duplication (dimer), allowing phase-shifted monomers to align across tandem repeat junctions. Sequences from repeat family A were queried against the database of repeats from family B using the settings -word_size 8 -reward 1 -penalty -1 -gapopen 2 -gapextend 2 -dust no, and we retained the best hit per query. As BLASTN reports local alignments, a global percentage identity was recovered by extending each alignment over the full query length, and assigning the 25% identity expected for random nucleotide sequence to unaligned regions^6^. The mean percentage identity across all within-pair alignments was used as the pairwise similarity estimate. Most recent common ancestor (MRCA) divergence times between all tip pairs were taken from the calibrated 325 species phylogeny. Following Melters et al^6^, pairwise similarity values were node-averaged by MRCA divergence time and fitted to an asymptotic exponential decay model:

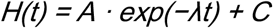

where H is node-averaged sequence identity, t is node age in millions of years, A is the decay amplitude, λ is the decay rate, and C is the empirical background similarity floor (estimated as a free parameter by non-linear least squares using scipy.optimize.curve_fit). We define the evolutionary half-life of the satellites as the divergence time at which similarity has decayed to halfway between its initial value and the empirical floor, which was calculated as t½=ln(2)/λ.

For satellite base composition comparisons, GC% content was calculated for all centromeric repeat family sequences. For each family, these values were compared to GC% from the set of proper chromosomes within a genome assembly, excluding unplaced scaffolds or contigs. For each satellite family, we tallied the copy number of repeats and created a ranked list. Zipf’s law states that the frequency of a value is inversely proportional to its rank in a sorted list of all values, such that the most frequent value appears approximately twice as often as the second most frequent, three times as often as the third, and so on. To test conformity to Zipf’s law we plotted the natural log (ln) of the repeat copy number rank, and the natural log of copy frequency, and tested for fit to a linear model.

Mean satellite similarity within and between chromosomes was calculated for each species by sampling 1,000 repeat pairs found on the same or different chromosomes, respectively, using an edit distance-based similarity metric, and performing paired t-tests to assess whether the similarities were significantly different. Similarities of holocentric repeats within arrays, between arrays on the same chromosome, and between arrays on different chromosomes, were calculated by sampling 1,500 arrays, or array pairs, for each scenario, and calculating a matrix of similarity values using 10 randomly sampled repeats from each array and calculating the mean similarity value for each matrix. Histograms of the data were plotted and paired t-tests were used to assess the significance of differences between the mean values. For sequence identity dot plots, we used either Re-Dotable (https://www.bioinformatics.babraham.ac.uk/projects/redotable/), or Moddotplot^88^.

### Satellite higher order repeat and regimentation detection

Satellite higher order repeats (HORs), understood as pairs of highly similar contiguous repeat regions, were identified using TRASH^46^. Each HOR consists of two blocks, where a block is a run of contiguous repeat monomers; the two blocks are paired such that each monomer in the first block is at least 96% similar to the corresponding monomer in the second block. A minimum block length of four monomers was specified, to increase detection specificity. To compare HOR abundance between genomes, each repeat monomer was assigned a HOR score. The score is the number of distinct partner monomers a repeat acquires in the second block of any HOR in which it appears in the first block, divided by the total number of repeats in the array (range 0-100%). To analyse HOR scores along the scaled length of centromeres, each array was divided into 30 length-scaled windows. Within each window, the summed HOR score was divided by the number of repeats in that window. Per-window values were then aggregated across plant, invertebrate and chordate species to give group-level profiles. For holocentric species with periodic satellite repeat arrays, all individual satellite arrays were used to generate scaled plots along holocentric arrays.

For visualisation of representative HOR patterns within satellite arrays, 50 consecutive repeat monomers from an array were extracted. For each window of repeats, all pairwise edit (Levenshtein) distances between the 50 monomer sequences were computed. Edit distances were converted to percentage sequence divergence by dividing by the mean monomer length. Divergence values greater than 20% were capped at this value to preserve visual contrast in the heatmap. To guide downstream clustering, the characteristic spacing between similar monomers was estimated directly from the divergence matrix. Monomer pairs falling in the lowest 5% of pairwise divergence values were identified, and the distance in monomers between these pairs was tabulated. If the most common such distance was 1, or no low-divergence pairs were found, no local period was inferred. Otherwise, a base periodic distance was estimated by iterative least-squares fitting. Each observed distance *d* was expressed as an integer multiple *k* of a candidate base distance *b* (k=round(d/b)), and *b* was updated as b = Σ(k·d)/Σ(k²) until convergence. This procedure is analogous in purpose to the harmonic-product step used for spectral period estimation in the regimentation analysis (the periodogram-based analysis of array periodicity) described below, but operates directly on the histogram of pairwise distances between highly similar monomers, rather than on a periodogram. Monomer sequences within each 50-repeat window were aligned with MAFFT^119^ (--auto). While edit distances are used for the pairwise-divergence display; MAFFT alignment-based distances are used for tree construction and clustering. Pairwise raw genetic distances were calculated from the alignment using ape and dist.dna (raw model, pairwise deletion), and a neighbor-joining tree was constructed using ape and njs. The same distance matrix was subjected to hierarchical clustering (average linkage), and monomers were partitioned into *k* clusters, with *k* set to the locally estimated period where available (default *k*=10). Each tree cluster was then assigned a unique colour from a rainbow palette. Membership of these tree clades is represented using matched colours along the x and y axis labels of the 2D divergence heatmap.

For each chromosome, the periodicity of HOR structures was assessed from the pairwise distances between HOR blocks, expressed in repeat-monomer units, rather than genomic coordinates. The raw counts of HOR block pairs were tabulated across all pairwise distances *d* and divided by the number of possible pairs at that distance (*N−d*, for an array of *N* repeats). This removes the combinatorial decay inherent to finite arrays, and yields a normalised signal that reflects biological HOR structure. The HOR distance signal was analysed using periodograms (detrended, demeaned, and using a 10% cosine taper) to estimate power as a function of frequency (i.e. cycles per repeat). The fundamental period was identified within a biologically plausible range (2-100 repeat units) using a harmonic product spectrum. Each candidate HOR frequency was scored by its mean log-power across itself and its first four harmonics, resolving ambiguity between a true fundamental repeat and the harmonics. Peak prominence (defined as power at the fundamental relative to local background) and spectral entropy (defined as the concentration versus spread of power across the full spectrum) were computed as auxiliary measures of periodic HOR signal strength. The significance of the strongest HOR period peak in the plausible range was tested with Fisher’s g-test, which is self-calibrated to the number of frequencies tested, and so allows for a consistent threshold (P≤0.001) across arrays and windows of different sizes. Arrays and windows were classified as ‘periodic’ or ‘heterogeneous’ accordingly.

The same procedure was applied within sliding windows (step=¼ window size) defined by the repeat index, to localize periodic and non-periodic HOR regions within each satellite array. Window size was set adaptively per chromosome (20× the array-wide estimated period, up to a default of 1,000 repeats, where the period was ≥50 repeats, or undetermined). Windows with fewer than 20 HOR block pairs were masked due to insufficient data. Per-window classifications were collapsed into contiguous regions. Short (≤3-window) heterogeneous HOR runs flanked by periodic HOR windows of matching period were reclassified as periodic, and treated as local dips in significance rather than true boundaries. Then, adjacent windows agreeing in HOR classification and period were merged, with overlapping boundaries split at their midpoint. For each chromosome, the fraction of array length occupied by each estimated HOR period (versus by heterogeneous HOR sequences) was calculated. This calculation summarizes the extent of periodic satellite higher order structure. Full implementation details are provided in available scripts (https://github.com/vlothec/eukaryotic_centromere_architecture).

To quantify how strongly centromere and genome metrics follow species phylogeny, we estimated Pagel’s λ^133^ across the time-calibrated 325-species chronogram. Pagel’s λ is a multiplier applied to the internal (off-diagonal) elements of the phylogenetic variance-covariance matrix, rescaling the shared ancestry expected under Brownian motion^134^. λ=0 removes all phylogenetic covariance with traits distributed as if on a star phylogeny, independent of relatedness. While λ=1 corresponds to trait evolution by Brownian motion along the tree. Genome size, chromosome number, satellite amount, and satellite monomer length, were log10-transformed prior to the analysis. λ was estimated by maximum likelihood with geiger::fitContinuous (model=“lambda”), and independently cross-validated with a generalised-least-squares fit (nlme::gls with an ape::corPagel correlation structure). Following Wilk’s intervals, we report profile-likelihood 95% confidence intervals, defined as the set of λ for which the likelihood-ratio statistic does not exceed the 95% quantile of the χ²₁ distribution, or equivalently all λ whose log-likelihood lies within 1.92 units of the maximum. Intervals were truncated at 0 or 1 where the profile did not fall by 1.92 before the boundary.

## Data availability

DToL genome assemblies are available at https://portal.darwintreeoflife.org, or at NCBI using the accession codes provided in **Supplementary Table 1**. DToL genome note citations for each assembly are also provided in **Supplementary Table 1**, where available. Code associated with data analysis is provided at the project GitHub page (https://github.com/vlothec/eukaryotic_centromere_architecture).

## Acknowledgements

We thank Liam Revell for advice in the ancestral state reconstruction modelling. We acknowledge grant support from DToL Wellcome Trust Grants 206194 and 218328 to MB; Wellcome Trust grants 218328 and 226458 to RD; UKRI/BBSRC grants BB/Y009487/1 and BB/V003984/1, ERC grant 101142254 EvoPanCen, and Human Frontier Science Program award RGP0025/2021 to IRH; Royal Society awards UF160222, RF/ERE/221032, URF/R/221024, RGF/R1/180006, RGF/EA/201030, and RF/ERE/210069 to AB; EMBO long-term postdoctoral fellowship ALTF224-2022 to RB; the Max Planck Society and ERC Starting Grant 101114879 HoloRECOMB and DFG Grant MA 870 9363/2-1 to AM; National Science Centre, Poland grant 2024/52/C/NZ2/00246 to PW; Foundation for Polish Science START fellowship for PW; a Postgraduate Fellowship for Studies Abroad from “la Caixa” Foundation LCF/BQ/EU24/12060051 to JGI; EPSRC-funded INFORMED-AI project EP/Y028732/1 to IK; GACR grant 24-10036S to JM; and computational resources and data storage facilities provided by the ELIXIR-CZ Research Infrastructure Project (LM2023055).

## Contributions

PW, EPR, JGI, MH, RD, AB, and IH designed the study. Genome annotation and analysis was performed by PW, EPR, JGI, MH, MZ, LO, KJ, RB, PS, MH, CZ, SK, MB, NG, PN, GF, FKT, MU-S, AM, JM, MB, RD, AB and IH. FISH and immunocytology experiments were performed by LO, LM, YMS, AM, and JM. PW, ER, MZ, LO, KJ, RB, PS, MH, GF, FKT, IAD, IK, JM, MB, RD, AB and IH wrote the paper, with input from all authors.

## Tree of Life Consortium author list

Mark Blaxter^1^, Nova Mieszkowska^2,3^, Neil Hall^4^, Peter Holland^5^, Richard Durbin^1,6^, Thomas Richards^5^, Matthew Berriman^1^, Paul Kersey^7^, Peter Hollingsworth^8^, Willie Wilson^2,9^, Alex Twyford^8,10^, Ester Gaya^7^ Mara Lawniczak^1^, Owen Lewis^5^, Gavin Broad^11^, Fergal Martin^12^, Michelle Hart^8^, Paul Flicek^12^, and Ian Barnes^11^

1 Wellcome Sanger Institute, Wellcome Genome Campus, Hinxton, Cambridgeshire CB10 1SA, UK

2 Marine Biological Association of the United Kingdom, Citadel Hill, Plymouth PL1 2PB, UK

3 University of Liverpool, Liverpool L69 3BX, UK

4 Earlham Institute, Norwich Research Park, Norwich NR4 7UZ, UK

5 Department of Biology, University of Oxford, Mansfield Road, Oxford OX1 3SZ, UK

6 Department of Genetics, University of Cambridge, Cambridge CB2 3EH, UK

7 Royal Botanic Gardens, Kew, Richmond, London TW9 3AE, UK

8 Royal Botanic Garden Edinburgh, Edinburgh EH3 5LR

9 University of Plymouth, Drake Circus, Plymouth PL4 8AA, UK

10 Institute of Ecology and Evolution, School of Biological Sciences, University of Edinburgh, Edinburgh EH9 3FL

11 Natural History Museum, Cromwell Road, London SW7 5BD, UK

12 EMBL-EBI, Wellcome Genome Campus, Hinxton, Cambridgeshire CB10 1SD, UK

